# A family-wide atlas of human connexin docking compatibility

**DOI:** 10.64898/2026.08.31.743583

**Authors:** Shan Ying, Joseph Peterman, Zheyan Song, Alissa T. Rzepski, Elizabeth Ransey

## Abstract

Gap junction (GJ) channels mediate direct intercellular communication by allowing the exchange of ions, metabolites, and signaling molecules between neighboring cells. Humans express 21 connexin (Cx) isoforms that can assemble into homotypic or heterotypic channels, creating a large potential interaction landscape that shapes tissue-specific communication networks. However, the rules governing which connexin isoforms can compatibly dock remain incompletely defined. Extracellular loop 2 (EL2) sequence features have been implicated in docking specificity and used to classify connexins into two canonical compatibility groups, K-N and H, but these assignments remain largely predictive. Most potential heterotypic connexin pairings have never been experimentally tested. This incomplete interaction map limits our ability to predict which connexin combinations can assemble, how isoform co-expression shapes intercellular communication, and how these relationships are altered or exploited in disease and engineered systems.

Here, we used the FETCH (Flow Enabled Tracking of Connexosomes in HEK Cells) assay to evaluate docking compatibility across the complete human connexin family. To support family-wide compatibility mapping, we used literature-supported heterotypic interactions to define a data-driven FETCH score threshold for high-confidence interaction compatibility. Homotypic FETCH measurements varied substantially across the 21 connexin isoforms, with 15 producing mean scores above the empirical threshold. We then extended FETCH analysis to all 210 pairwise heterotypic isoform combinations. The resulting interaction landscape largely recapitulated expected motif-class relationships, including enrichment within the two canonical compatibility groups, but also identified neighboring-group interactions and unexpected cross-group pairings that represented clear exceptions to class-based predictions. Consistent with these findings, pairwise EL2 motif similarity was only modestly associated with threshold-based interaction classification, indicating that EL2 similarity alone was insufficient to predict compatibility outcomes. Together, these findings suggest that motif class provides a broad organizing framework for connexin compatibility, but that pairwise docking specificity also depends on yet-unresolved isoform-specific determinants that produce neighboring-group relationships and clear cross-group exceptions. Notably, Cx46, a lens Cx also associated with melanoma and breast cancers, emerged as a broadly permissive isoform capable of interacting with partners from both major compatibility groups and more than half of the connexin family. Together, these findings establish the first family-wide experimental atlas of human connexin docking compatibility, defining canonical interactions, previously unrecognized pairings, and exceptions to established compatibility rules. This atlas provides a foundation for defining the molecular determinants of connexin specificity, understanding how isoform diversity shapes intercellular communication, and designing gap junction channels with controlled docking behavior.

## Introduction

Gap junction (GJ) channels mediate the direct cytoplasmic exchange of ions, metabolites, and signaling molecules^1^, including ATP^2^, cAMP^3^ and miRNAs^4^, between neighboring cells, enabling coordinated activity across tissues such as the heart, brain, and epithelium^5^. GJ channels are formed when connexin (Cx) hemichannels on adjacent cells dock across the extracellular space to create intercellular pores (Fig. 1A). The human genome encodes 21 Cx isoforms^6–8^ that share a conserved membrane topology (Fig. 1B) but differ in sequence features that affect permeability, gating, trafficking, oligomerization, and docking compatibility. Most tissues co-express multiple Cx isoforms, and this diversity allows cells to build specialized communication networks with distinct molecular composition and signaling properties^5, 9, 10^. Consistent with these broad physiological roles, mutations and aberrant expression of Cx proteins are associated with numerous human diseases including congenital sensorineural deafness^11, 12^, developmental disorders^13^, epileptic conditions^14,15^, neurodegeneration^16^ and cancer^17^.

**Figure 1.**
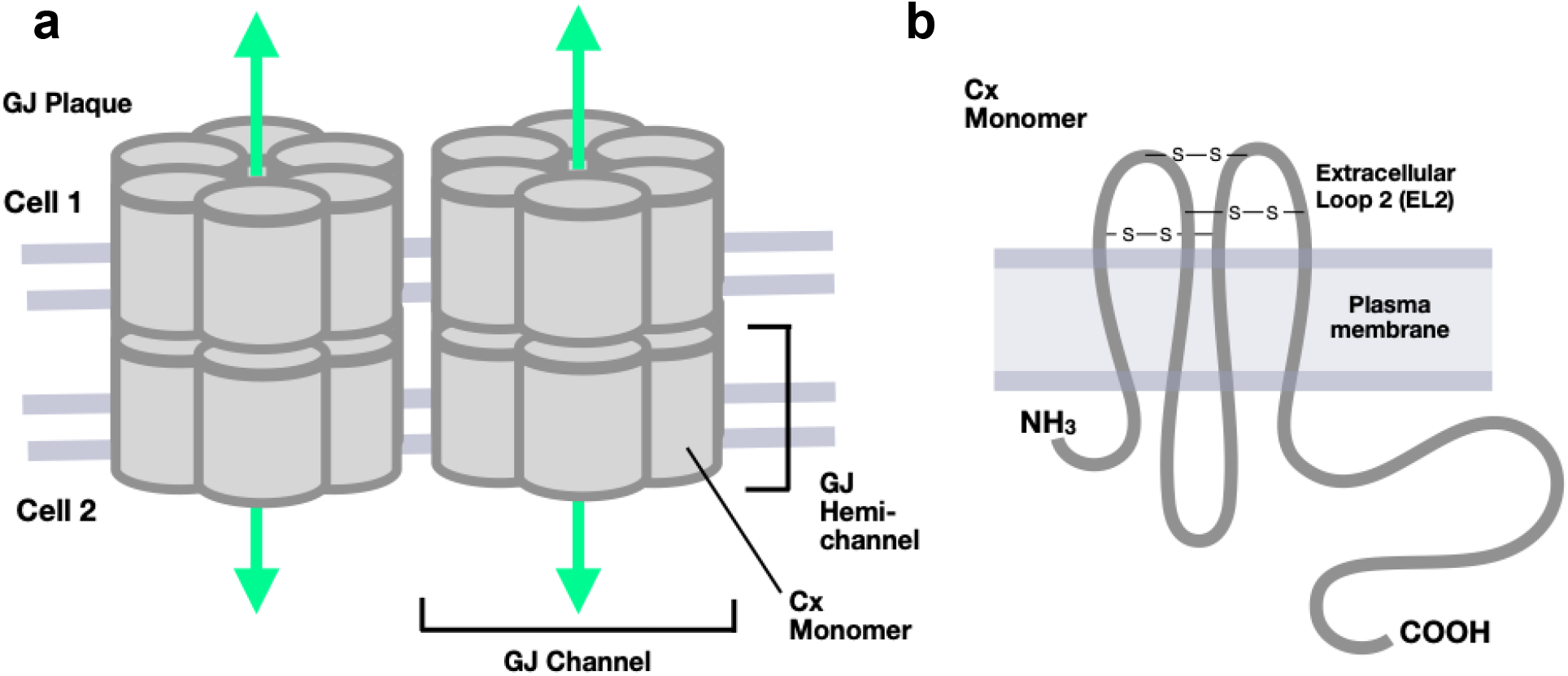
Composition of Gap Junction Intercellular Channels. **(A)** Schematic representation of a gap junction (GJ) plaque connecting two neighboring cells. Individual connexin monomers assemble into hexameric hemichannels within each cell membrane. Docking of compatible hemichannels across the extracellular space generates complete intercellular GJ channels, multiple of which cluster together to form larger junctional plaques. A representative GJ channel within the plaque is highlighted. **(B)** Schematic topology of a connexin monomer showing the cytoplasmic N- and C-termini, four transmembrane domains (TM1–TM4), intracellular loop (CL), and extracellular loops (EL1 and EL2). The extracellular loop 2 (EL2) domain, a major determinant of hemichannel docking specificity and compatibility, is highlighted.

A central unresolved question is how Cx isoform identity determines which hemichannels can compatibly dock to form stable gap junction channels. The potential interaction landscape across the human connexin family is large and complex, because each connexin can form homotypic (same type) channels and heterotypic (two-type) channels with a subset of other isoforms^18^. Heterotypic docking is particularly interesting because channels assembled from two different connexins can exhibit properties that differ from either corresponding homotypic channel, including altered conductance, voltage-dependent gating, permeability, molecular selectivity, or directionality of intercellular transfer^19–21^. At the same time, docking compatibility does not necessarily establish that a resulting channel supports robust or physiologically equivalent communication. Some pairings may form morphologically stable junctions yet display weak, asymmetric, or context-dependent function, whereas others may create communication pathways between cell populations that would otherwise remain uncoupled. Defining heterotypic compatibility is therefore an essential first step towards understanding the broader interaction space available to co-expressed connexins: which neighboring cells can directly communicate, which channel combinations can form, and how might these unique channels expand, restrict, or qualitatively alter intercellular signaling in native and disease contexts.

Despite the biological importance of heterotypic interactions, the molecular rules that determine which isoforms can dock remain incompletely understood. Prior work has implicated extracellular loop sequences, particularly extracellular loop 2 (EL2) (Fig. 1B), as key determinants of docking specificity. These findings have led connexins to be organized into compatibility groups based on conserved EL2 motif architectures, including the canonical K-N (Group 1; Φ(K/R)CxxxPCPNxVDCΩΨS) and H (Group 2; ΦxCxxxPCPHxVDCΩΨS) classes, with heterotypic compatibility generally expected to occur preferentially among isoforms within the same class^18, 22^. Although this framework provides a useful basis for predicting compatible connexin pairings, it remains largely inferential, as most heterotypic combinations across the human connexin family have never been experimentally tested.

A comprehensive experimental atlas is therefore needed to define which connexin pairs support docking compatibility; to evaluate the limits of existing predictive frameworks; and to identify candidate interactions for functional testing. Such a map would also support protein engineering efforts aimed at designing connexins with desired docking specificity for programmed cell–cell communication. Indeed, our recent development of a synthetic connexin pair with selective docking specificity demonstrated that engineered connexins can function in the mammalian nervous system to enable targeted circuit synchronization and behavioral interrogation^23^.

Here, we show that the FETCH (Flow Enabled Tracking of Connexosomes in HEK Cells) assay^23, 24^ can be used to quantify homotypic and heterotypic interaction compatibility across all 21 human connexin isoforms. We first measured homotypic pairings to assess how each isoform behaves in the assay, identifying connexins that produce robust docking-associated GJ internalization as well as isoforms with comparatively low homotypic FETCH activity. We measured heterotypic FETCH scores across all pairwise isoform combinations (n = 210) and used literature-supported compatibility assignments to establish a data-driven threshold for high-confidence interactions. The resulting interaction landscape largely aligned with expected compatibility relationships based on motif-class assignment, including enrichment within canonical compatibility groups, but also revealed neighboring and distant cross-group interactions that extended beyond class-based predictions. Additionally, we found that EL2 motif similarity was only weakly correlated with FETCH scores, indicating that motif similarity alone is insufficient to predict the full interaction landscape captured by the assay. Finally, we identified Cx46 as a broadly compatible bridging isoform, capable of forming stable interactions with partners from both major compatibility groups.

Overall, this work generates a family-wide experimental atlas of human connexin docking compatibility, defining canonical interactions, previously unrecognized pairings, and cross-boundary relationships that extend beyond motif-class predictions. By organizing these interactions into a systematic compatibility map, this atlas tests the limits of existing predictive frameworks and identifies candidate heterotypic relationships for further mechanistic and functional evaluation. Because docking compatibility alone does not establish channel conductance, permeability, or physiological function, these interactions should be viewed as experimentally defined compatibility potential rather than evidence that all detected pairs support equivalent intercellular communication. This resource therefore provides a foundation for future studies of connexin interaction specificity and for investigating how isoform-specific differences shape gap junction assembly, remodeling, and communication in native and engineered contexts.

## Results

### Homotypic FETCH captures broad but variable docking-associated transfer across human connexins

During the GJ life cycle, connexin monomers assemble into hexameric hemichannels, traffic to the plasma membrane, dock with compatible hemichannels on apposed cells, and cluster into junctional plaques containing tens to thousands of intercellular channels^1^ (Fig. 1A). These plaques, or portions of them, are subsequently internalized into one of the two coupled cells through a coordinated clathrin-dependent^25, 26^ endocytic–exocytic process resulting in internalized double-membrane vesicular structures termed annular gap junctions^27^ or connexosomes^5^. Because connexosomes arise directly from pre-existing docked junctional plaques, their formation provides a downstream readout of successful and sufficiently stable GJ docking. Connexosome generation remains incompletely understood and has been explicitly characterized for only a subset of gap junctions, including those generated by Cx43 and Cx26. However, a recent study reported internalization of junctional structures formed by several additional connexin isoforms, including some that lack canonical endocytic motifs^28^. Recently, we developed FETCH, a high-throughput assay that uses dual-color connexin signal as a readout of docking-associated connexin transfer and internalization (Fig. 2A)^23, 24^. To determine whether FETCH could support an all-isoform interaction atlas, we first tested whether the assay detected docking-associated connexin transfer across all 21 homotypic pairings, where most human connexins are expected to form gap junctions.

**Figure 2.**
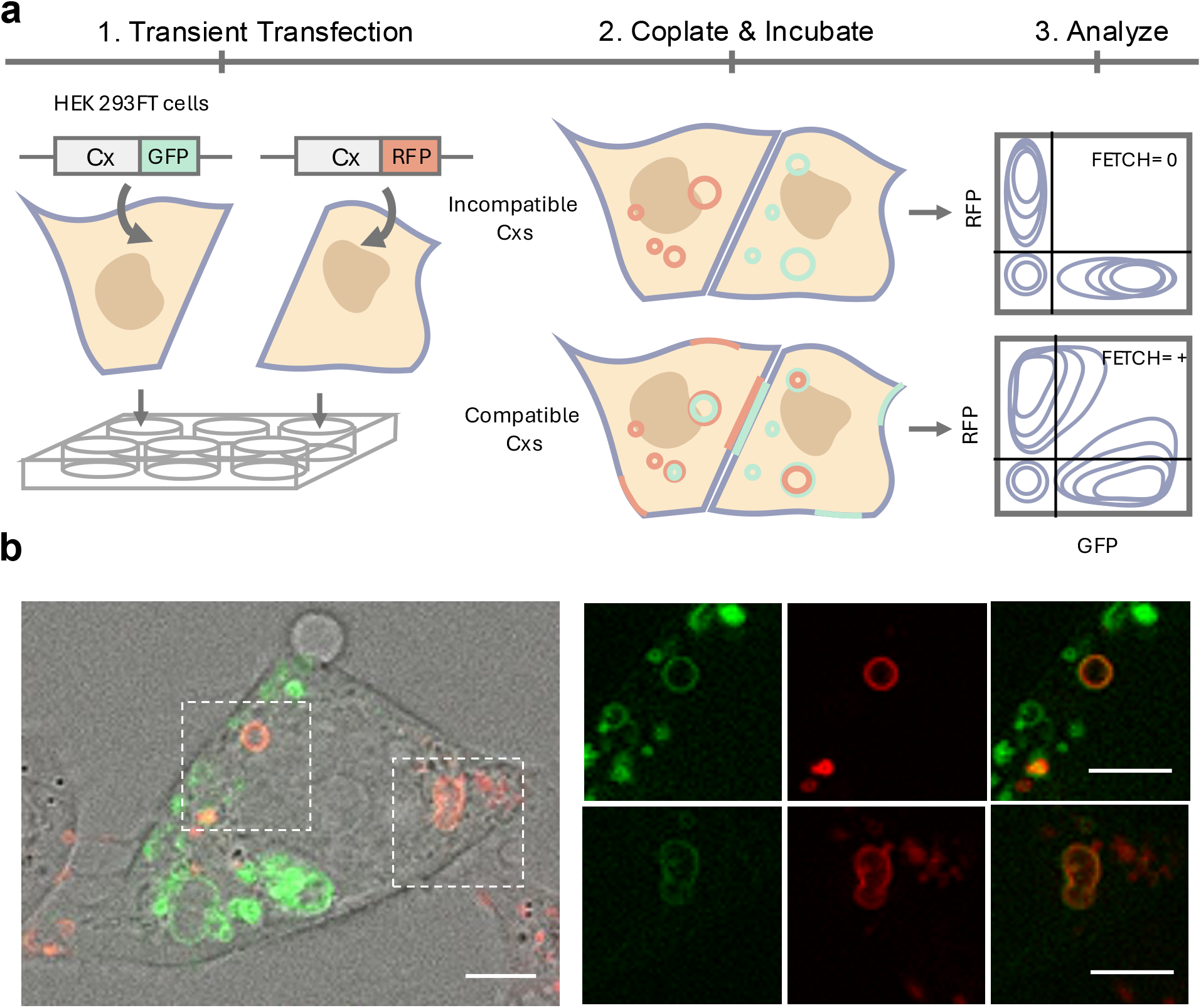
Connexosomes can distinguish Cx docking compatibility. **(A)** Schematic overview of the compatibility readout. Separate HEK293FT cell populations are transiently transfected with connexin constructs tagged with spectrally distinct fluorescent proteins. Following co-plating and incubation, incompatible connexin pairs are expected to produce limited docking-associated transfer and low dual-color signal by flow cytometry. In contrast, compatible connexin pairs can form docked gap junctions and generate internalized dual-color connexin-positive structures, producing increased dual-color signal and higher FETCH scores. Mock flow cytometry plots illustrate the expected readout for incompatible and compatible connexin pairs. **(B)** Representative fluorescence image of an individual cell from a Cx43-expressing co-culture showing internalized gap junction structures.. Insets show dual-color connexin-positive internalized structures in merged and single-channel views. Scale bars are 5 *μ*m.

In the FETCH workflow, separate HEK293FT cell populations are transiently transfected with spectrally distinct fluorescently-tagged connexin constructs, such as Cx-mEmerald and Cx-mCherry, and allowed to express overnight (Fig. 2A, left). The two populations are then combined at equal proportions and co-plated in wells of the same size used for the initial transfections. Co-plated cells are then incubated overnight, during which time docked GJs can form and undergo remodeling and internalization. Cells expressing compatible connexins are expected to dock efficiently and form gap junctions and connexosomes, whereas incompatible pairings are expected to exhibit little or no stable docking and correspondingly limited junction and connexosome formation (Fig. 2A, center). Finally, cells are trypsinized, fixed, and analyzed by flow cytometry, where dual-positive events provide a quantitative readout of connexosomes generated by two cell populations (Fig. 2A, right). Representative imaging of homotypic Cx43 co-cultures after trypsinization showed dual-color connexin-positive internalized structures supporting the ability of FETCH to detect docking-associated connexin transfer and internalization (Fig. 2B). For analysis of the FETCH data, a flow cytometry gating strategy was designed to sequentially identify and select the HEK293FT cell population, singlets, and fluorescent protein-positive cells (Fig. S1). Within the fluorescent protein-positive population, cell events are displayed on a two-parameter fluorescence plot, with mEmerald and mCherry defining the two axes. Quadrant gates are established using kernel density estimation of the untransfected population to distinguish negative, single-positive, and dual-positive events. To standardize comparisons across connexin pairs, the FETCH score is calculated as the fraction of dual-fluorescent color positive events relative to all fluorescent events, correcting for variation in transfection efficiency: FETCH score = Q2/(Q1 + Q2 + Q3) (Fig. S1).

In preparation for the all-isoform homotypic FETCH analysis, the full set of human connexin constructs was transiently transfected into HEK293FT cells and evaluated by high-resolution confocal imaging for subcellular localization (Fig. 3), as well as by western blotting and widefield imaging for assessment of overall expression and relative transfection efficiency. (Fig. S2). The imaging analyses confirmed construct expression and provided visual context for general localization patterns consistent with the connexin family. Western blot signal intensity and apparent banding patterns varied across isoforms, consistent with isoform-specific differences in expression, trafficking, detergent sensitivity, and biochemical behavior. These experiments were intended to confirm construct detectability rather than optimize conditions for each isoform.

**Figure 3.**
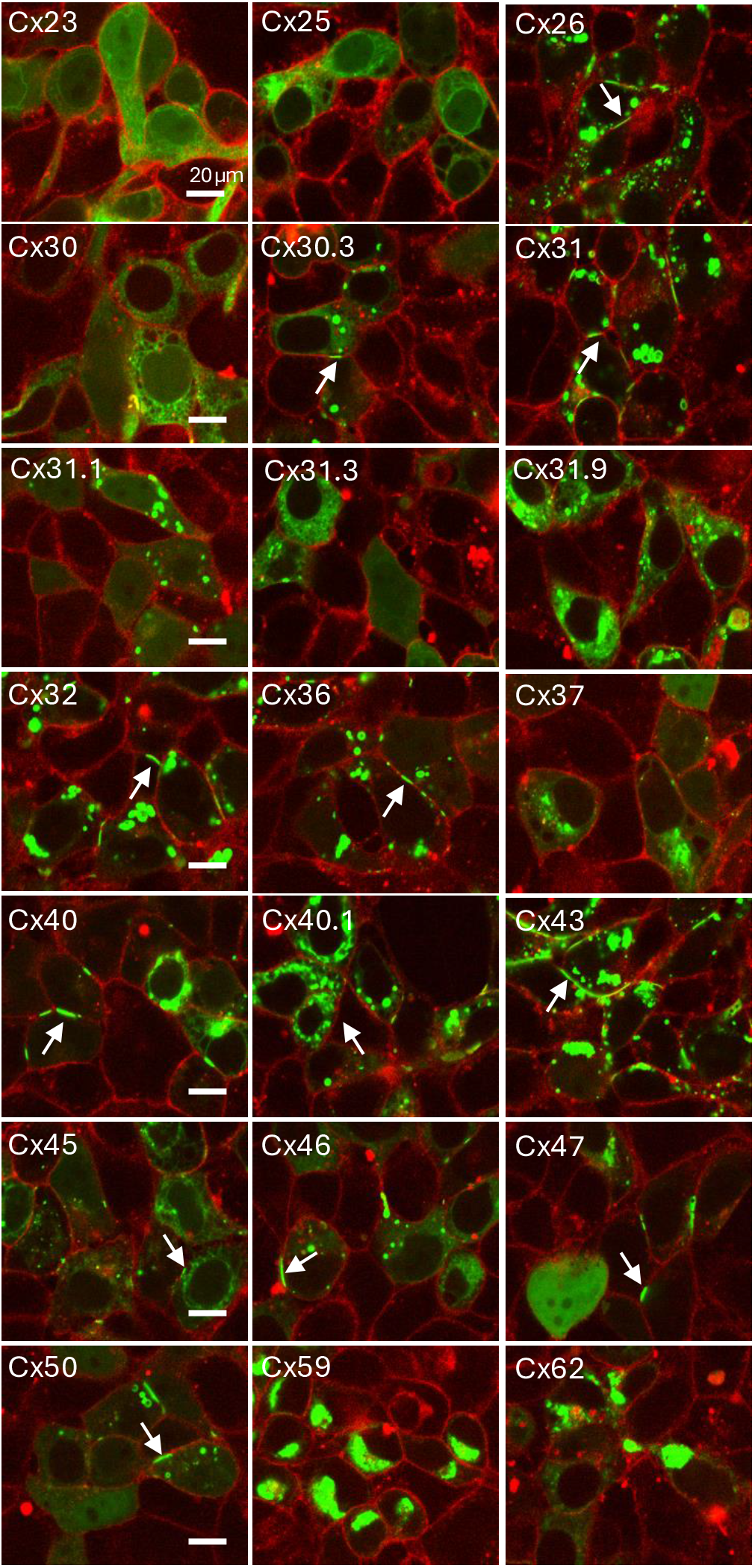
Human connexin isoforms exhibit diverse subcellular localization and plaque morphologies. Representative fluorescence images of HEK293FT cells expressing each of the 21 human connexin isoforms tagged with fluorescent protein. Images were collected under shared imaging conditions to provide a visual survey of connexin localization and intracellular distribution across the human connexin family. The panel highlights the diversity of isoform-specific expression patterns, including differences in junctional enrichment, intracellular localization, and apparent plaques (white arrows), providing context for interpreting isoform-dependent differences in FETCH scores and compatibility profiles.

Before comparing FETCH profiles across the connexin family, we first established a consistent sequence-based framework for organizing isoform identity. We aligned the extracellular loop 2 (EL2) sequences of all 21 human connexins and classified each isoform according to conserved motif architecture. This analysis incorporated previously defined K-N and H class assignments and extended classification across the remaining isoforms, assigning connexins to canonical K-N or H classes, related K-N-like or H-like groups, or an “Other” category^18^ for sequences that did not clearly conform to these motifs (Fig. S3). These classifications were used to color-code subsequent FETCH plots and to compare interaction patterns across the connexin family. With this classification framework in place, we first examined the ability of FETCH to capture homotypic docking across the full human connexin family. As nearly all connexin isoforms are expected to form homotypic gap junctions, homotypic pairings provided an initial test of isoform-specific FETCH performance. Using the complete construct set, we quantified homotypic FETCH scores for all 21 isoforms. Experiments were performed as previously described, with cytoplasmic fluorescent proteins included as nonjunctional controls. Each homotypic pairing was evaluated across at least three biological replicates, defined as independent transfections, with at least four technical replicate samples per experiment. Replicate homotypic FETCH scores were plotted as distributions for each connexin and ranked by mean value, revealing substantial isoform-dependent variation in flow cytometry profiles (Fig. S4) and FETCH score readout (Fig. 4).

**Figure 4.**
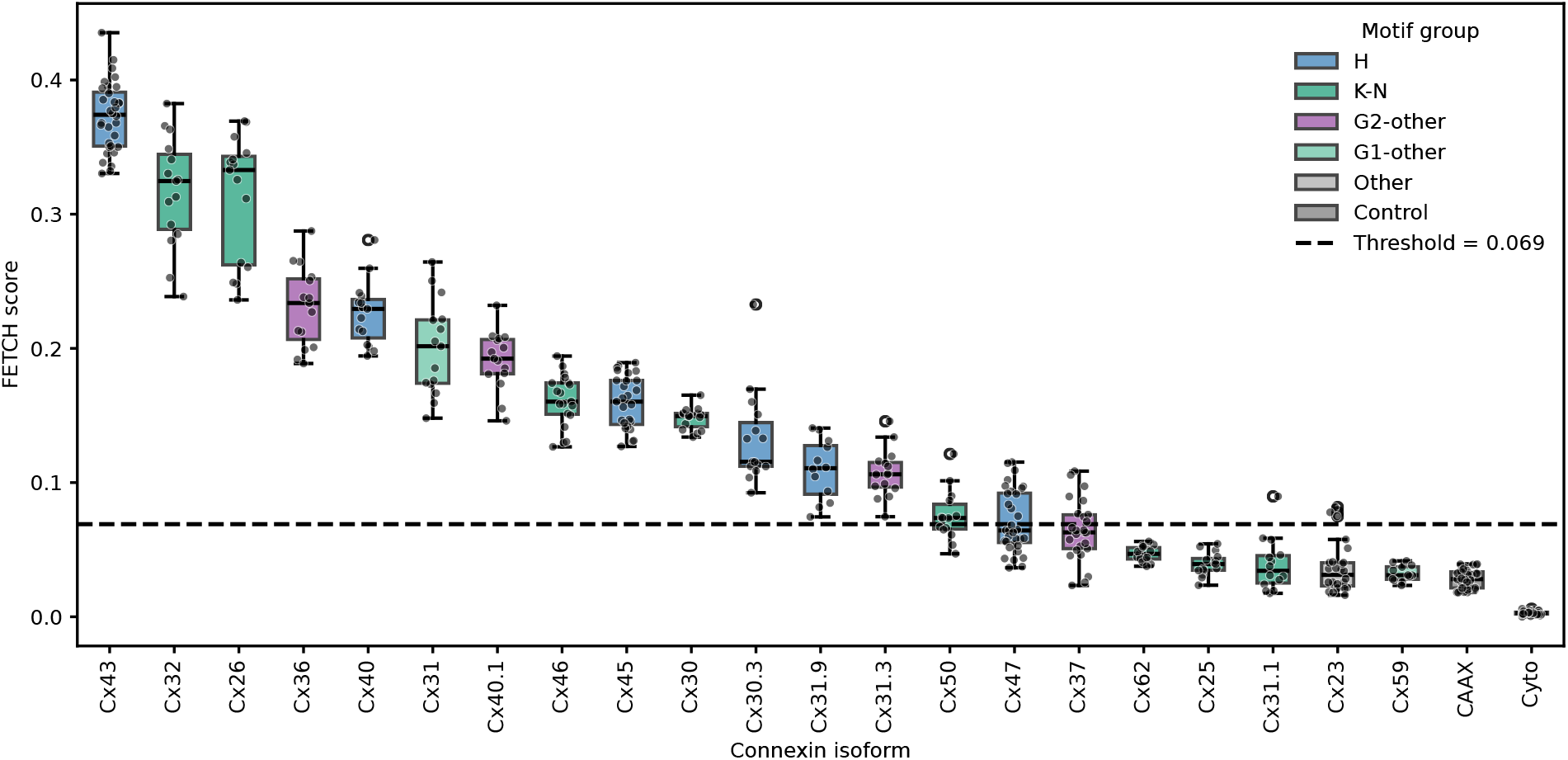
Homotypic FETCH interaction scores for all 21 isoforms. Homotypic FETCH scores were quantified for all 21 human connexin isoforms and plotted as replicate-level distributions. Isoforms are displayed in descending order by mean homotypic FETCH score. Each box represents the distribution of replicate homotypic FETCH scores for one isoform, with the center line indicating the median and the box indicating the interquartile range. The dashed line indicates the reference value used in subsequent heterotypic analyses to classify high-confidence interactions, 0.069, and is shown here for visual comparison rather than as an independently optimized homotypic cutoff.

The highest mean homotypic FETCH scores were observed for Cx43, Cx32, and Cx26, at 0.373, 0.317, and 0.312, respectively. In contrast, Cx62, Cx25, Cx31.1, Cx23, and Cx59 produced among the lowest mean scores, ranging from 0.047 to 0.032. To contextualize homotypic FETCH activity relative to the heterotypic screen, we plotted homotypic scores against the empirical threshold used to classify high-confidence heterotypic interactions. This threshold was derived from the heterotypic dataset by comparing FETCH scores with literature-supported compatibility assignments and applying ROC analysis with Youden’s J statistic to identify the score that best separated reported compatible from incompatible pairs. The resulting threshold was 0.069.

By mean homotypic FETCH score, 15 of 21 isoforms exceeded this value, consistent with detectable homotypic docking-associated transfer for most human connexins. Six isoforms fell below this value: Cx23, Cx25, Cx31.1, Cx37, Cx59, and Cx62. Several of these have prior evidence of limited or uncertain homotypic channel formation, including Cx23^29, 30^ and Cx31.1^31^, while Cx25 and Cx59 remain poorly characterized and Cx62 may be affected by proteolytic processing^32^. Cx37 is a notable exception, as it is known to form functional homotypic gap junctions^33^ and its performance in subsequent heterotypic FETCH screening suggests that its slightly subthreshold score may reflect threshold or assay-specific behavior instead of poor junction forming capacity. Because the threshold is derived from heterotypic benchmarking, it is used here as a contextual reference rather than a strict homotypic cutoff. Together, these measurements define isoform-specific assay behavior that can inform the interpretation of their respective heterotypic FETCH profiles.

Finally, because HEK293FT cells may express endogenous connexins that could contribute to FETCH scores, primarily Cx43 and Cx45^33^, we asked whether homotypic FETCH scores were maintained in cells lacking these endogenous isoforms. We therefore repeated the homotypic FETCH measurements in Cx43/Cx45 double-knockout (DKO) HEK293 cells and compared isoform-level scores between the HEK293FT and DKO backgrounds. Although the two cell lines were generally similar in morphology and growth behavior, we observed modestly smaller cell size, slower growth and lower transfection efficiency in the Cx43/Cx45 DKO cells than in HEK293FT cells. It was unclear whether this reflected reduced expression, increased cell death, or both, but lower fluorescence limited our ability to reliably collect FETCH data for some isoforms. Thus, for Cx59, duplicate sequential transfection increased the proportion of fluorescently expressing cells and enabled reliable homotypic FETCH measurement.

Overall, homotypic FETCH profiles were broadly similar between HEK293FT and DKO cells. Strong Pearson and Spearman correlations indicated preservation of both score magnitude and isoform rank order across cell backgrounds (Pearson r = 0.85, p = 1.29 x 10^-6^; Spearman ρ = 0.85, p = 1.03 x 10^-6^ Fig. S5). Threshold-based classifications were also highly concordant: all 15 isoforms classified as homotypic FETCH-positive in HEK293FT cells remained above threshold in the DKO background, while 6 subthreshold isoforms remained below threshold in both backgrounds. The only discordant cases were Cx23 and Cx31.3, two isoforms previously reported to exhibit weak or nonfunctional homotypic channel behavior^30, 34^. In both cases, the differences represented borderline threshold shifts: from 0.036 (Cx23, negative) and 0.107 (Cx31.3, positive), respectively, in HEK293FT cells to 0.076 (Cx23, positive) and 0.051 (Cx31.3, negative) in DKO cells. Together, these results indicate that endogenous Cx43 and Cx45 do not substantially alter homotypic FETCH classifications under these assay conditions, supporting the use of HEK293FT cells for heterotypic compatibility mapping.

### Family-wide heterotypic FETCH analysis identifies canonical, atypical, and previously unreported connexin compatibility relationships

After defining homotypic FETCH profiles for individual connexin isoforms, we performed combinatorial heterotypic FETCH experiments across all pairwise human connexin combinations (n = 210). Heterotypic assays were conducted similarly to homotypic assays, with the exception that replicate samples were distributed across reciprocal fluorescent-protein configurations, such that each connexin partner in each pair was tested with each FP tag. This design minimized FP-specific signal bias and generated quantitative FETCH measurements for every possible heterotypic pair. Replicate measurements across reciprocal fluorescent-protein configurations were averaged to obtain a single mean FETCH score for each heterotypic pair, and the resulting values were assembled into a complete quantitative interaction matrix.

To identify broader compatibility patterns, we first examined the heterotypic interaction profile of each isoform individually and then organized and ordered the global matrix according to similarity among these profiles (Supplemental Table 1, Fig. 5A). Specifically, pairwise similarity between continuous FETCH interaction vectors was calculated using correlation distance, followed by hierarchical clustering with average linkage. The resulting dendrogram order was applied to the heterotypic interaction matrix, revealing clusters of isoforms with similar interaction behavior. This organization revealed two prominent motif-class-associated clusters, with K-N motif connexins (e.g., Cx26, Cx32 and Cx46) and H motif connexins (e.g., Cx43, Cx45 and Cx47) grouping into largely distinct regions of the matrix. This separation indicates that connexins with shared EL2 motif architecture tend to exhibit similar heterotypic FETCH profiles, consistent with established compatibility group relationships. A third, low-scoring cluster included isoforms with no known heterotypic compatibility (i.e., Cx36)^18^, and also isoforms which exhibited comparatively weak homotypic FETCH activity (i.e., Cx23, Cx25, Cx31.1, Cx59 and Cx62). More broadly, the heterotypic interaction landscape was sparse, with most isoform pairs producing only weak FETCH signals (< 0.10). In contrast, several isoforms exhibited elevated FETCH scores with partners outside their canonical compatibility group.

**Figure 5.**
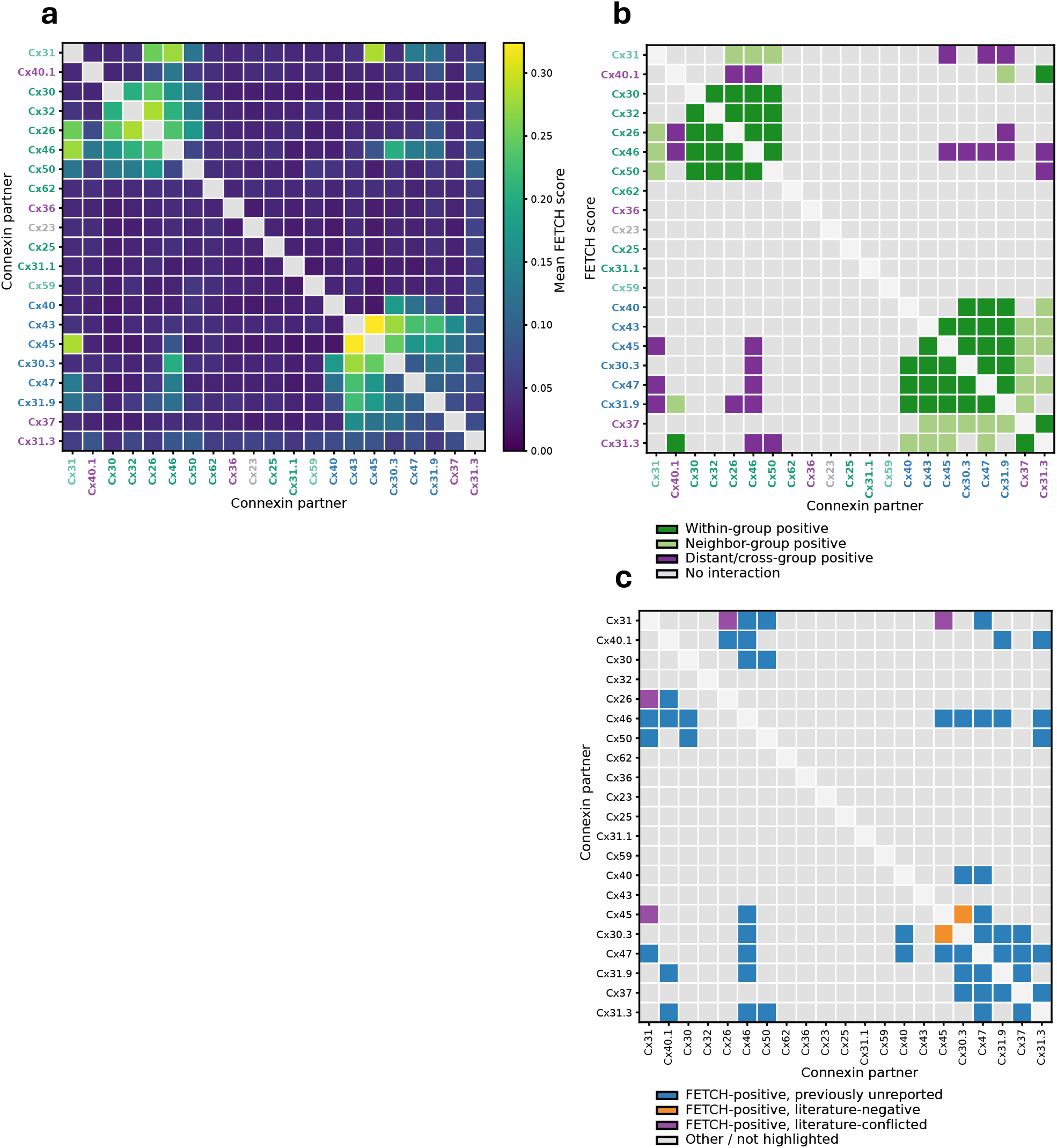
Global organization of heterotypic Cx compatibility relationships. **(A)** Continuous heterotypic FETCH interaction matrix across 21 human connexin isoforms. Connexins are ordered by unsupervised hierarchical clustering based on similarity of their complete heterotypic FETCH interaction profiles. Color intensity reflects experimentally measured FETCH scores for each heterotypic pairwise interaction. The clustered compatibility landscape reveals structured compatibility neighborhoods together with progressive overlap between canonical compatibility groups. **(B)** Thresholded heterotypic interaction matrix generated using an empirical FETCH interaction threshold of 0.069, determined by ROC analysis and Youden’s J statistic relative to previously reported connexin compatibility classifications. Isoforms are displayed in the same clustered order as in panel A. Dark green indicates high-confidence within-group interactions, light green indicates neighboring-group interactions, purple indicates distant/cross-group interactions, and gray indicates interactions below the threshold. **(C)** Literature-status matrix comparing thresholded FETCH-positive heterotypic interactions with published compatibility reports. Positive interactions were classified as previously supported, newly identified with no prior compatibility report, or literature-discordant, where FETCH detected a high-confidence interaction despite previous reports of incompatibility or negative functional coupling. Of the 50 high-confidence FETCH-positive interactions, 26 represented newly identified compatibility relationships and 1 represented literature-discordant positive interactions.

To establish a threshold for defining high-confidence positive interactions, we benchmarked heterotypic FETCH scores against literature-supported compatibility assignments from previously reported heterotypic connexin studies. Of the 210 possible heterotypic isoform combinations, literature-based compatibility information was available for 62 pairs, including 28 reported compatible interactions and 22 reported incompatible or nonfunctional pairings (Supplemental Table 2; Fig. S6A)^21, 31, 35–88^. Pairs with conflicting interaction reports across studies or assay types (n=12) were assigned to a separate conflicted literature category and excluded from threshold calculations. Using these curated assignments, we performed ROC analysis and applied Youden’s J statistic to identify the FETCH score that best separated literature-supported compatible and incompatible pairs. This analysis identified 0.069 as the threshold that maximized concordance between FETCH-based classification and literature-supported compatibility (Fig. S6B). Applying this threshold to the full heterotypic dataset identified 50 high-confidence positive interactions among 210 evaluated pairs which we organized into a thresholded matrix (Fig. 5B). Of these threshold positive interactions, 25 occurred within compatibility groups, 13 occurred between neighboring groups (e.g., H to H-like), and 12 occurred between distant or cross-group pairings (H to K-N).

Analysis of the thresholded matrix showed that cross-boundary interactions were largely concentrated between related compatibility classes. H-group connexins frequently paired with H-like connexins (G2-other), whereas K-N-group connexins were enriched for compatibility with K-N-like connexins (G1-other; Fig. 5B). Nevertheless, we also observed distant and cross-group interactions that represented direct exceptions to canonical compatibility categories. Several isoforms, including Cx31, Cx46, and Cx31.3, contributed disproportionately to these cross-boundary relationships, with high-confidence interactions spanning multiple motif classes. In particular, Cx46 formed several bona fide K-N-to-H pairings that would not be predicted based on motif class, including interactions with Cx45 (0.09), Cx47 (0.14), Cx30.3 (0.20), and Cx31.9 (0.12).

To evaluate how FETCH-defined positive interactions compared with prior studies, we placed the thresholded interaction matrix in the context of curated literature reports. Among the 50 threshold-crossing pairs, 26 lacked prior compatibility reports and therefore represent newly determined potential Cx interactions, whereas only 1 was a literature-discordant interaction in which FETCH detected compatibility despite previous negative reports^71^ (i.e. Cx30.3/Cx45; 0.25) and 2 FETCH-identified positive interactions have conflicted reports in the literature^37, 42^ (Cx26/Cx31, 0.25 and Cx31/Cx45, 0.28; Fig. 5C). Together, these results demonstrate that heterotypic connexin compatibility is broadly organized by canonical motif class but extends beyond strict class boundaries. By integrating FETCH-based thresholding with literature context, this analysis expands the experimentally supported human connexin compatibility landscape by defining high-confidence canonical and atypical interactions, identifying newly recognized pairs, resolving conflicted cases, and highlighting literature-discordant interactions for future orthogonal validation.

### Pairwise EL2 motif similarity is modestly associated with threshold-positive heterotypic FETCH interactions

The presence of several interactions that crossed canonical compatibility-group boundaries suggested that motif-class assignment alone may be too coarse a metric to fully predict heterotypic FETCH behavior. Although the K-N and H classes are defined by conservation of selected EL2 sequence features, individual isoforms within each class retain substantial sequence diversity. We therefore asked whether pairwise EL2 sequence similarity was associated with FETCH score across the full heterotypic dataset.

To test the relationship between EL2 motif sequence and FETCH score directly, we plotted pairwise EL2 motif similarity, calculated as percent identity across the aligned reported-length EL2 motif region (17 amino acids; shown in Fig. S3), with heterotypic FETCH scores in HEK 293FT cells across all evaluated heterotypic pairs. This comparison revealed broad dispersion in FETCH scores across the range of EL2 similarities, with most distant/cross-group pairs falling below the empirical interaction threshold and a smaller set of threshold-positive pairs emerging despite divergent EL2 motifs (Fig. 6). Reflecting this broad scatter, EL2 motif sequence similarity showed only a weak positive association with FETCH score, with Spearman ρ = 0.19 and Pearson r = 0.26. Thus, greater EL2 motif sequence similarity was associated with slightly higher FETCH scores overall, but motif similarity alone was not a strong predictor of quantitative FETCH signal.

**Figure 6.**
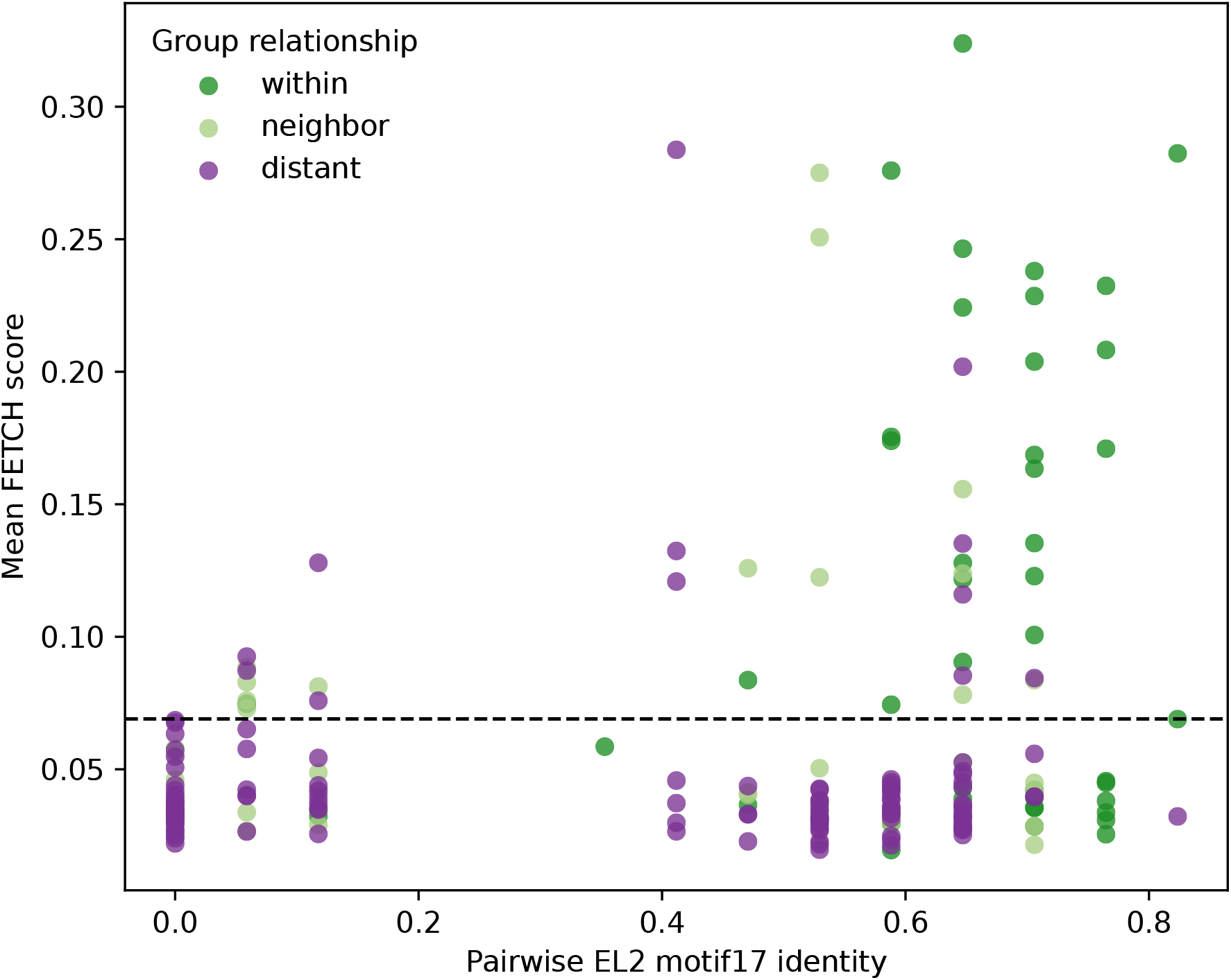
EL2 motif similarity is associated with heterotypic compatibility but does not fully predict FETCH score. Pairwise EL2 motif similarity plotted against experimentally measured heterotypic FETCH scores for all tested connexin pairs. Pairwise motif similarity was calculated across the reported-length EL2 compatibility motif window using global alignment-based percent identity. EL2 motif similarity exhibited statistically detectable but modest associations with heterotypic FETCH behavior (Spearman ρ = 0.17, Pearson r = 0.22), indicating that EL2 motif architecture contributes to, but does not fully determine, heterotypic compatibility relationships.

Quantitative FETCH score magnitude is influenced by multiple isoform-specific factors, including expression, trafficking, membrane availability, docking stability, and internalization. Given these constraints, we asked whether pairwise EL2 motif similarity was associated with quantitative FETCH score and whether it more effectively distinguished threshold-positive from threshold-negative interactions. Using the empirically defined FETCH threshold of 0.069, threshold-positive pairs exhibited greater reported-length EL2 motif similarity than threshold-negative pairs, with median identities of 0.647 and 0.529, respectively (Mann–Whitney *P* = 1.32 × 10⁻⁴). However, EL2 sequence similarity alone provided only modest discrimination between threshold-positive and threshold-negative interactions (AUC = 0.678). Logistic regression likewise indicated that increasing EL2 motif identity was associated with greater odds of a FETCH-positive interaction classification (odds ratio = 1.225 per 10 percentage-point increase in identity, 95%CI = 1.074-1.397, p = 0.0025)

We next asked whether this limited predictive performance reflected the use of the complete EL2 motif rather than localized sequence features. To address this, we compared exact amino acid identity with physicochemical similarity based on conservative substitutions across the full 17-residue motif and overlapping 12-and 7-residue sliding windows (Supplemental Table 3).Restricting the analysis to shorter sequence windows produced only modest improvement. The strongest localized signal was observed for exact identity across residues 4–10, increasing the AUC from 0.678 to 0.705 and the Spearman correlation with continuous FETCH score from ρ = 0.189 to ρ = 0.228. However, no individual window strongly separated FETCH-positive from FETCH-negative interactions or more accurately tracked interaction magnitude. Likewise, allowing conservative substitutions with similar physicochemical properties did not improve predictive performance relative to exact amino acid identity, yielding a full-motif AUC of 0.650 and a maximum windowed AUC of approximately 0.69. Together, these analyses indicate that EL2 sequence conservation captures broad patterns of docking compatibility that largely align with the established connexin compatibility groups, while also revealing pair-specific exceptions that are not fully resolved by localized sequence identity or physicochemical conservation alone.

To test unexpected compatibility relationships identified in the EL2 similarity analysis, we focused on connexin pairs with relatively high FETCH scores despite low EL2 motif sequence similarity. These pairs were readily identifiable in the scatterplot as above-threshold distant-group interactions (purple points; Fig. 6). We ranked these heterotypic pairings by FETCH score and selected representative 8 representative examples (Cx31/Cx45, Cx46/Cx30.3, Cx47/Cx46, Cx31.9/Cx31, Cx46/Cx40.1, Cx46/Cx31.9, Cx47/Cx31 and Cx45/Cx46) for two-color, high-resolution imaging. Following sequential transfection, we assessed whether each pair formed heterotypic gap junctions (plaques). All assayed pairs formed two-color gap junction plaques, providing independent support that the unexpected cross-group interactions detected by FETCH assemble into junctional structures (Fig. S7, white arrows).

Together, these analyses indicate that EL2 motif architecture captures broad patterns of connexin docking compatibility, with interactions enriched within canonical or related compatibility classes. However, the substantial dispersion in FETCH scores and the presence of validated distant-group interactions show that motif classification alone does not fully predict individual pairwise outcomes. These exceptions suggest that docking specificity is shaped not only by broad EL2 class identity, but also by sequence-level features, structural context, and additional isoform-specific determinants that refine compatibility beyond simple motif-based expectations.

### Cx46 emerges as a highly connected isoform in the heterotypic compatibility atlas

Having defined the global heterotypic compatibility landscape and examined its relationship to EL2 sequence features, we next asked whether individual isoforms exhibited unusually broad or atypical interaction profiles. This analysis identified Cx46 as a high-connectivity outlier in the atlas. Cx46 exhibited the largest number of threshold-positive heterotypic interactions of any connexin in the dataset, with 11 high-confidence heterotypic partners. It also showed the greatest number of distant or cross-group interactions, with 6 partners outside its expected compatibility class (Cx30.3, Cx47, Cx40.1, Cx31.9, Cx45 and Cx31.3; Fig. 5B), and accounted for 8 previously unreported FETCH-positive interactions (Cx31.9, Cx47, Cx30.3, Cx45, Cx40.1, Cx30, Cx31 and Cx31.3; Fig. 5C). Thus, Cx46 represented one of the clearest examples of an isoform whose compatibility profile extended beyond its motif-class prediction.

Given the breadth of unexpected Cx46 interactions identified in the heterotypic screen, and the knowledge that Cx46 can hetero-oligomerize with Cx43^89^, we asked whether this broad interaction profile could reflect contributions from endogenous connexins – Cx43 and Cx45. To test this, we transiently transfected Cx46 into Cx43/Cx45 DKO HEK cell line and compared its localization and heterotypic FETCH activity across backgrounds with those observed in the HEK 293FT cells. Representative images showed that Cx46 formed GJ plaques in both cell types, consistent with Cx46 trafficking and plaque localization being preserved in the absence of endogenous Cx43 and Cx45 (Fig. 7A). In agreement with the all-isoform, homotypic 293FT cell to DKO cell comparison (Fig. S5), Cx46 heterotypic FETCH scores were strongly preserved between HEK 293FT and DKO cells, with Pearson and Spearman correlation analyses indicating similar score magnitude and partner rank order across backgrounds (Pearson r = 0.96, p = 5.87 x 10^-11^; Spearman ρ = 0.85, p = 1.65 x 10^-6^; Fig. 7B). Pair-level differences were small (Fig. S8A) and largely fell within replicate-variability-derived equivalence bounds (Fig. S8B), supporting the conclusion that the broad Cx46 profile was not substantially driven by endogenous Cx43 or Cx45. These data increase confidence that the Cx46 interactions detected in the atlas reflect compatibility relationships between the transfected isoform pairs.

**Figure 7.**
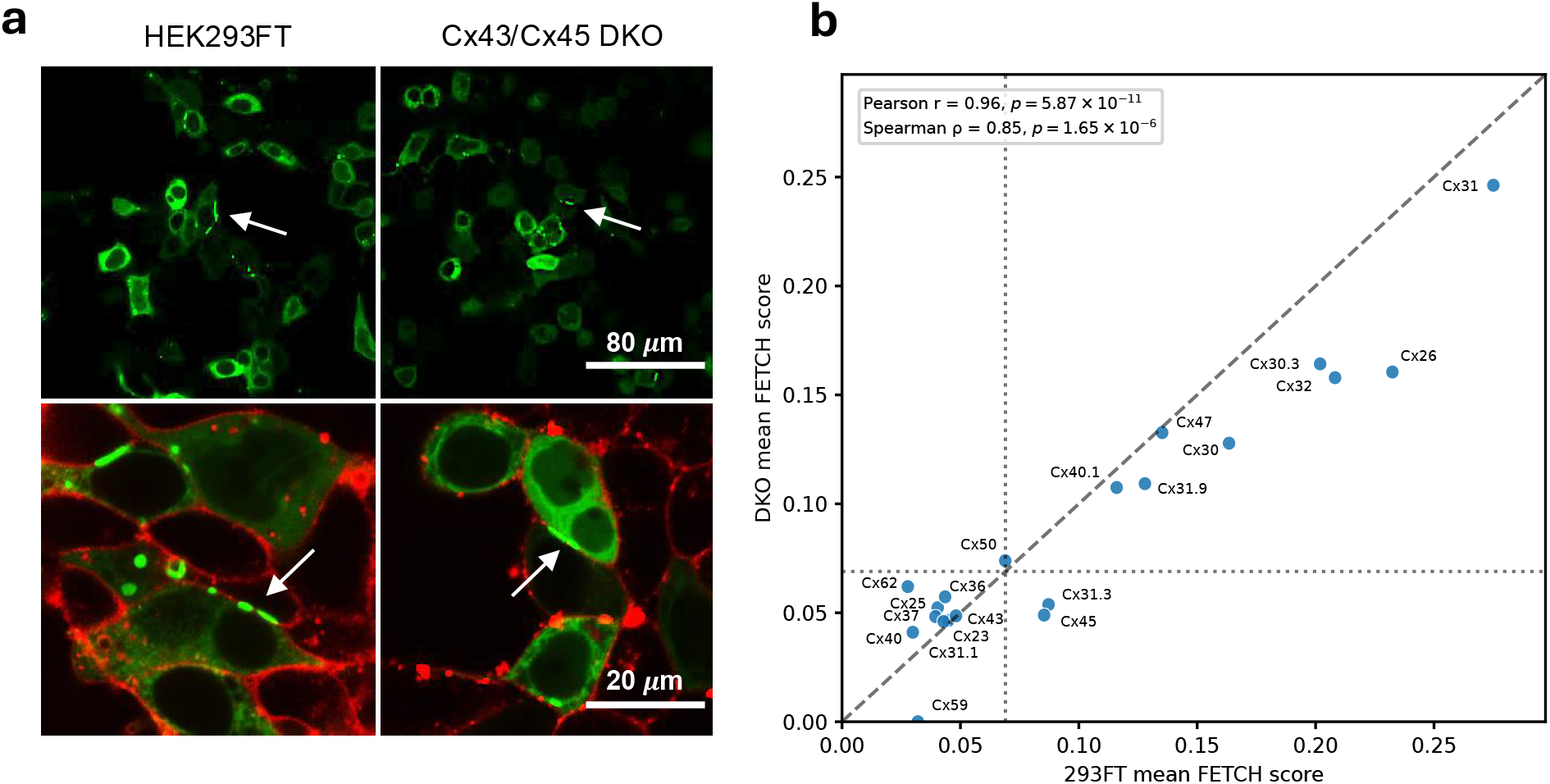
Cx46 exhibits broad heterotypic compatibility that is preserved in Cx43/Cx45 double-knockout cells. **(A)** Representative images of fluorescently tagged Cx46 expressed in parental HEK293FT cells and Cx43/Cx45 double-knockout cells. Images provide visual context for Cx46 expression and localization in the two cellular backgrounds used for FETCH analysis. White arrows indicate gap junction plaques. **(B)** Comparison of Cx46-containing heterotypic FETCH scores measured in parental HEK293FT cells and Cx43/Cx45 double-knockout cells. Each point represents one matched Cx46 heterotypic pair. Scores were strongly concordant between cell backgrounds, supporting the conclusion that the broad Cx46 compatibility profile was not substantially driven by endogenous Cx43 or Cx45.

To interpret Cx46 heterotypic interactions in the context of isoform-specific FETCH variability, we next compared each Cx46 heterotypic FETCH score with the corresponding partner’s homotypic activity (Fig. S9). Eleven isoforms exhibited both strong homotypic activity and above-threshold heterotypic interactions with Cx46, supporting their classification as high-confidence positive partners. In contrast, Cx36, Cx40, and Cx43 showed robust homotypic activity but remained below the heterotypic threshold with Cx46, consistent with true pair-specific incompatibility. Six additional isoforms (Cx23, Cx25, Cx31.1, Cx37, Cx59, and Cx62) scored weakly in both homotypic and heterotypic measurements and therefore warrant greater scrutiny, as their low Cx46 scores may reflect limited assay performance rather than definitive incompatibility. This comparison supports the broad compatibility of Cx46 while distinguishing likely positive and negative interactions from pairs that require additional validation.

Consistent with the global heterotypic interaction analysis, comparing Cx46 heterotypic FETCH scores with EL2 motif similarity revealed that Cx46 partner preferences were not fully explained by compatibility class or nearest-neighbor sequence relationships. Although Cx46 showed the highest reported-length EL2 motif sequence identity to Cx50 (82.4%), consistent with their shared K-N motif classification, the Cx46–Cx50 pair produced only a modest FETCH score among Cx46-positive interactions, ranking 11th of 11 (0.069, Table 1). Conversely, Cx46 showed high-confidence interactions with several neighboring and distant cross-class partners, including Cx31, Cx30.3, Cx47, Cx40.1, Cx31.9, Cx45, and Cx31.3, even though their EL2 sequence similarities ranged from 5.9% to 64.7%. This pattern distinguishes Cx46 as a broadly compatible isoform whose strongest partners include several interactions outside its nearest EL2 sequence neighbors.

**Table 1.**
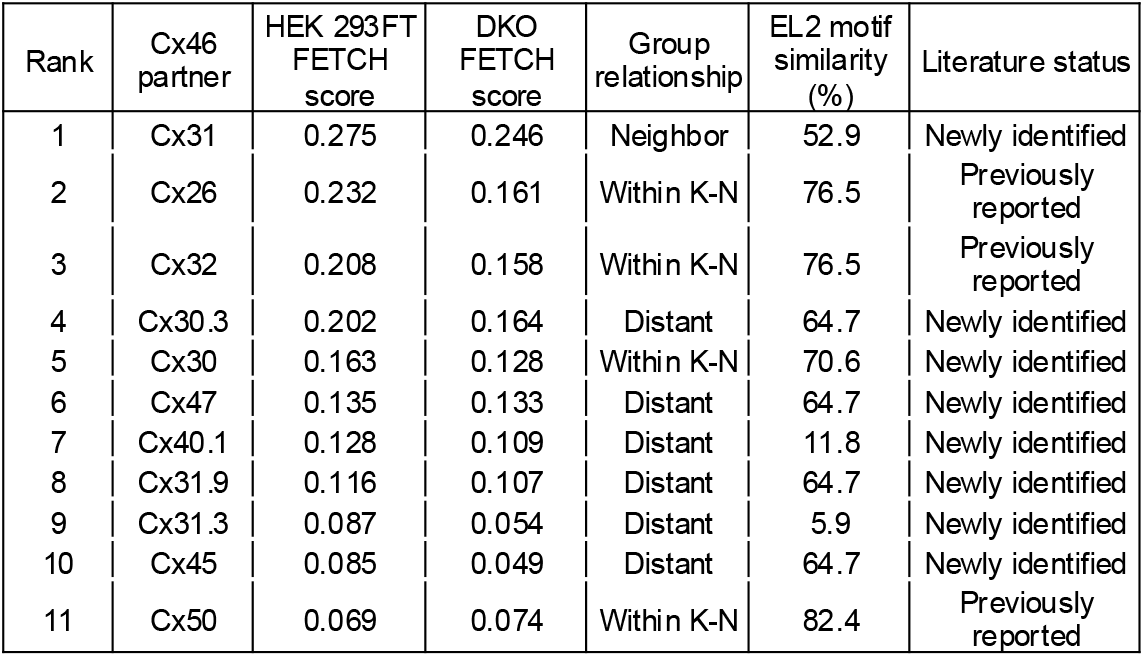
Cx46 heterotypic partner profile. Table summarizing Cx46-containing heterotypic interactions that exceeded the empirically defined FETCH threshold. Interactions are ranked by mean FETCH score and annotated by compatibility-group relationship, pairwise EL2 motif similarity, and literature status.

| Rank | Cx46 partner | HEK 293FT<br>FETCH<br>score | DKO<br>FETCH<br>score | Group<br>relationship | EL2 motif<br>similarity<br>(%) | Literature status |
| --- | --- | --- | --- | --- | --- | --- |
| 1 | Cx31 | 0.275 | 0.246 | Neighbor | 52.9 | Newly identified |
| 2 | Cx26 | 0.232 | 0.161 | Within K-N | 76.5 | Previously reported |
| 3 | Cx32 | 0.208 | 0.158 | Within K-N | 76.5 | Previously reported |
| 4 | Cx30.3 | 0.202 | 0.164 | Distant | 64.7 | Newly identified |
| 5 | Cx30 | 0.163 | 0.128 | Within K-N | 70.6 | Newly identified |
| 6 | Cx47 | 0.135 | 0.133 | Distant | 64.7 | Newly identified |
| 7 | Cx40.1 | 0.128 | 0.109 | Distant | 11.8 | Newly identified |
| 8 | Cx31.9 | 0.116 | 0.107 | Distant | 64.7 | Newly identified |
| 9 | Cx31.3 | 0.087 | 0.054 | Distant | 5.9 | Newly identified |
| 10 | Cx45 | 0.085 | 0.049 | Distant | 64.7 | Newly identified |
| 11 | Cx50 | 0.069 | 0.074 | Within K-N | 82.4 | Previously reported |

The broad Cx46 interaction profile is particularly interesting given the physiological and pathological contexts in which Cx46 has been implicated. Cx46 is best known for its role in the lens, where mutations are associated with cataracts^90, 91^, but it has also been linked to multiple cancer settings, including glioblastoma cancer stem cells^92^, melanoma^93^, and breast cancer^94–96^ In these tissues, Cx46 is likely expressed alongside distinct repertoires of endogenous connexins. Its expanded heterotypic compatibility profile therefore raises the possibility that Cx46 function may depend not only on its expression level, but also on which other connexins are present in each cellular context. These findings identify Cx46-containing heterotypic pairs as strong candidates for future validation and suggest that isoform-specific compatibility may contribute to tissue-and disease-specific functions of Cx46.

## Discussion

This study establishes a family-wide FETCH-based atlas of human connexin docking compatibility. To our knowledge, it represents the first systematic evaluation of all 21 homotypic and 210 heterotypic human connexin pairings using a single standardized assay. By measuring FETCH scores across all 21 human connexin isoforms, we transformed a fragmented body of prior pairwise observations into a comprehensive interaction map, generating empirical evidence for more than 150 previously untested interactions and identifying multiple high-confidence, previously unreported heterotypic pairings. This resource allowed us to evaluate how docking-associated interaction patterns align with canonical EL2 motif-based compatibility classes, prior reports, and pairwise EL2 motif sequence comparisons, while also revealing atypical cross-class relationships that expand the known human connexin interaction landscape.

The homotypic interaction dataset revealed substantial isoform-to-isoform variation in FETCH detectability across the connexin family, providing an internal reference of isoform-specific assay performance. Most connexins produced strong homotypic signal, supporting broad applicability of FETCH across the human connexin family. However, several isoforms produced comparatively weak signal even when paired with themselves, potentially reflecting limited homotypic channel formation, as reported for Cx23 and Cx31.1, or isoform-specific differences in plaque formation, trafficking, membrane availability, or internalization. The homotypic FETCH score variability helped contextualize heterotypic results, as low FETCH scores may reflect selective incompatibility or broader isoform-specific limits on assay detection.

Comparison of FETCH results between HEK293FT cells and Cx43/Cx45 double-knockout cells provided important insight into the influence of endogenous connexins on the assay. Homotypic classifications across all isoforms, as well as selected heterotypic classifications involving Cx46, were largely preserved after removal of endogenous Cx43 and Cx45. This supports the conclusion that measured FETCH interactions primarily reflected the transfected connexin pairs rather than background endogenous connexin interference. Practically, these findings suggest that connexin-null or engineered background cell lines may be valuable for targeted validation experiments, but that HEK293FTs (a commercially available cell line) can be used effectively for systematic compatibility screening in FETCH studies.

The global heterotypic interaction matrix generated by FETCH revealed both expected and unexpected patterns of connexin compatibility. Benchmarking against literature-supported compatibility assignments enabled the definition of a data-driven threshold for high-confidence heterotypic interactions, identifying 50 positive interactions among the 210 evaluated pairs. While many of these interactions occurred within canonical compatibility groups (n=25/50) or between neighboring motif classes (n=13/50), others spanned distant or cross-group (n=12/50) relationships that are not readily predicted by existing class-based frameworks. Placing these threshold-positive interactions in the context of prior reports further highlighted the value of a family-wide experimental map. Many high-confidence FETCH-positive interactions lacked prior experimental compatibility evidence (n=26/50), while one interaction conflicted with previous reports of incompatibility or absent functional coupling and two others helped reconcile contradictory findings in the literature. Although these newly identified and literature-discordant interactions do not, by themselves, establish channel function, they expand the set of candidates connexin pairs capable of supporting docking-associated transfer and internalization in the FETCH assay.

Though observed compatibility relationships largely favored existing canonical classifications, pairwise EL2 motif similarity was only weakly associated with quantitative FETCH score. This weak score-level relationship is consistent with FETCH acting as an integrated readout influenced by isoform-specific expression, trafficking, and post-docking processing. We therefore asked whether EL2 similarity was more informative for distinguishing FETCH-positive from FETCH-negative interactions using the empirical interaction threshold. Using this threshold-based classification, greater EL2 similarity was associated with FETCH positivity, providing moderate discrimination between FETCH-positive and FETCH-negative interactions and corresponding to increased odds of a positive interaction. These associations were not substantially improved by focusing on smaller sequence windows or by allowing conservative amino acid substitutions. These findings suggest that EL2 sequence features help organize the compatibility landscape, while unexpectedly compatible pairs may reflect isoform-specific sequence differences not fully captured by broad motif-class assignments and interaction patterns not predicted by existing structural and biochemical models. Although we focused on EL2 because of its established role in docking specificity, the limited predictive value of EL2 similarity alone suggests that additional extracellular or structural features may help refine pairwise compatibility. EL1 is one possible feature to examine in future studies, although its reported conservation across canonical compatibility groups^22^ makes it less likely to serve as a significant determinant of class-level specificity.

Cx46 emerged as the most highly connected isoform in the heterotypic FETCH atlas, with several novel and cross-group interactions identified. Consistent with the global heterotypic analysis, the Cx46 profile was not well predicted by EL2 motif similarity, as its closest EL2 motif match was its weakest FETCH partner and several lower-similarity partners produced robust FETCH scores. These findings identify Cx46, a connexin associated with numerous cancers, as a particularly interesting candidate for future studies aimed at determining how docking compatibility, channel function, and tissue-specific connexin co-expression combine to shape isoform-specific gap junction biology.

This study provides a family-wide experimental map of human connexin docking compatibility and positions FETCH as a scalable, accessible framework for systematic connexin interaction testing. Although all 21 connexins were evaluated, confidence in individual interaction identifications varies with homotypic FETCH performance. Most connexins produced robust homotypic signal, increasing confidence in their heterotypic measurements, whereas pairs involving weak homotypic performers should be interpreted more cautiously. FETCH detects connexin-dependent fluorescent signal transfer between cells following co-culture. Positive interactions are therefore consistent with gap junction docking and internalization but do not independently establish channel conductance or definitive ultrastructure. As an integrated readout influenced by expression, trafficking, junction formation, and internalization, FETCH may not distinguish true incompatibility from limited assay detectability in low-scoring pairs. This distinction is especially important for atypical or literature-discordant interactions, which may represent biologically meaningful compatibility relationships, assay-sensitive interactions, or cases in which docking and channel function are separable. Likewise, the empirical interaction threshold is a literature-calibrated classification tool rather than an absolute biological boundary, and continuous FETCH scores are retained to preserve differences in interaction magnitude.

As with any overexpression-based assay, FETCH may identify interactions favored by elevated connexin abundance or simplified cellular context. However, the structured organization of the matrix, overall agreement with prior reports, and low scores observed for most heterotypic pairs argue against nonspecific overexpression-driven transfer as the primary explanation for the observed compatibility landscape. Interactions identified in this in vitro system may also not occur in native tissues, where expression, localization, cell type, and isoform abundance constrain which connexins can interact. Nevertheless, these findings define pairwise compatibility potential and provide a foundation for testing its functional, physiological, and engineering relevance.

Altogether, these findings reveal an organized landscape of human connexin docking compatibility that extends beyond canonical motif-class boundaries and simple sequence-based expectations. EL2 class strongly shapes this landscape, but isoform-specific sequence variation, broader extracellular or structural features, and cellular context likely further refine pairwise interaction specificity. Although FETCH-positive interactions do not by themselves establish channel conductance or native-tissue occurrence, they identify pairwise compatibility potential that can now be tested through functional validation, targeted mutagenesis, and physiological models. By defining canonical interactions, atypical cross-class relationships, and previously unrecognized connexin pairs, this atlas provides a foundation for structure-guided modeling and machine learning approaches aimed at defining the molecular rules governing connexin pairing, gap junction assembly, and engineered intercellular communication.

## Materials and Methods

### Construct cloning and preparation

Connexin gene information was procured from the National Center for Biotechnology Information (NCBI, ncbi.nlm.nih.gov) and the Ensembl genome browser (ensembl.org). The human codon-optimized genes were ordered from Integrated DNA Technology (IDT, idtdna.com). Genes were initially subcloned into mEmerald-N1 (addgene:53976) vectors using In-Fusion cloning (takarabio.com), resulting in carboxy-terminal fluorescent protein fusion constructs. To generate mCherry-labeled counterpart connexin constructs, the mCherry2 gene (gifted from Yongxin Zhao, Carnegie Mellon University) was subcloned in as an mEmerald replacement.

### Cell Lines

HEK 293FT cells were purchased from Thermo Fisher Scientific (cat# R70007). The Cx43/Cx45 double-knockout HEK293-derived tsA201 cell line, a gift from Steve Reichow (Oregon Health & Science University), was generated commercially by GenScript using CRISPR–Cas9 targeting of GJA1 and GJC1. Isogenic clones were isolated by limiting dilution and validated as full allelic knockouts by PCR and Sanger sequencing. All delivered cell populations tested negative for mycoplasma.

### Cell Culture

Both HEK cell lines were grown according to Thermo Fisher Scientific instructions: cultures were grown in 10 cm tissue culture treated dishes in high-glucose DMEM (Thermo Fisher Scientific, cat#11965092) supplemented with 10% Fetal bovine Serum (FBS), 6mM L-glutamine, 0.1 mM MEM non-essential amino acids and 1mM MEM sodium pyruvate in a 5% CO2, 37°C incubator. Cells were passaged via trypsinization every 2-3 days at ∼60-80% confluency.

### Transient Transfection

HEK 293FT cells were plated onto 10 *μ*g/ml Fibronectin coated multi-well dishes to achieve ∼75% confluency after overnight incubation. For transfection, HEK 293FT cells were transfected with polyethylenimine (PEI, Polysciences, cat# 23966), DKO HEK cells were transfected with Lipofectamine™ 3000 (Thermo Fisher Scientific, cat# L3000015) both according to manufacturer’s instructions. For PEI, DNA was diluted in Opti-MEM with PEI in a 1:3 ratio (*μ*g of DNA: *μ*L of reagent) and incubated at room temperature for 10 minutes. For Lipofectamine™ 3000: DNA was diluted in Opti-MEM with P3000™ in a 1:2 ratio (*μ*g of DNA: *μ*L of reagent) and subsequently mixed with Lipofectamine™ reagent and incubated at room temperature for 10 minutes. Following incubation, PEI-DNA or Lipofectamine-DNA complexes were added dropwise to wells of plated cells. Treated cells were then incubated at 37°C for 18-24 hrs, followed by media change. Expression of the connexin-FP constructs were evaluated at 18-24 hrs post transfection via widefield or confocal microscopy and western blotting.

### Image Acquisition

<u>For low-resolution, widefield</u> expression and localization imaging, HEK cells were plated onto 10 *μ*g/ml Fibronectin coated multi-well plastic plates. Cells were transiently transfected as described above and imaged at ∼20-24 hrs post transfection. Images were acquired on a Keyence BZ-X810 imaging system using a 40W LED light source, conventional fluorescence filters and a 10X, Plan Fluor, NA: 0.30 Ph1 objective. <u>For high-resolution confocal</u> <u>imaging</u>, cells were plated onto 10 *μ*g/ml Fibronectin coated 35 mm, glass-bottom Mattek dishes (Mattek, P35GC-1.5-14-C) and imaged with a Zeiss LSM 880 inverted confocal microscope using Argon/2, HeNe 594nm and HeNe633nm lasers, conventional fluorescence filters and a 63X, Zeiss PL Apo, NA: 1.4, Oil Immersion Objective, Images were taken with 1024 x 1024-pixel resolution. Membrane staining was achieved using Biotium live cell stain (CellBrite® Fix 555, cat# 30088), which was used according to manufacturer’s instructions.

### For imaging two-color plaques

Cells were plated onto 10 *μ*g/ml Fibronectin coated 35 mm, glass-bottom Mattek dishes and transfected as previously described but transfected twice, sequentially: first with the Cx-mEmerald construct and 4 hrs later with the Cx-mCherry construct. This allows for two separate populations to be transfected in one well. <u>Connexosome Imaging:</u> Cells were prepared using the same general workflow as the FETCH assay. Separate cell populations were plated, transfected, combined, replated into multi-well plastic plates, and incubated overnight. Two hours before imaging, co-plated cells were trypsinized and replated onto 35-mm glass-bottom MatTek dishes coated with 10 *μ*g/mL fibronectin. Images were acquired in filtered mode on a Keyence BZ-X810 imaging system using a 40 W LED light source, conventional fluorescence filter sets, and a 100× Plan Apo 1.45 NA oil-immersion objective

### Cell Lysate preparation and Western Blotting

For protein analysis via western blotting, transfected cells were washed twice with room temperature PBS, trypsinized, collected and centrifuged at 3,000 x g for 5 minutes at 4°C. Pellets were lysed by the addition of 10 mM Tris, pH 7.5, 1 mM EDTA, 1% SDS, 1X protease and phosphatase inhibitor cocktail (Cell Signaling Technology, cat# #5872), 20 U/mL DNase I (NEB, M0303S) and sonicated. Lysates were quantified via detergent compatible Bradford assay (Thermo Fisher Scientific, cat# 23246). For visualization, samples were separated using gel electrophoresis on 4-20% SDS-Page gels. Separated proteins were transferred to PVDF membrane in transfer buffer (25 mM Tris, 190 mM glycine, 20% methanol, final pH 8.3), for 45 min at 110V. Subsequently, membranes were blocked in 5% milk for 1 hr, followed by 3 hr or overnight at 4°C incubation with primary antibody (1:1000) in 5% milk with shaking. Blots were then washed three times with TBST (20 mM Tris, pH 7.5, 150 mM NaCl and 0.1% Tween 20) for 5 minutes and incubated with Horse radish peroxidase (HRP) conjugated secondary antibody (1:5000), followed by three washes with TBST for 5 minutes. Blots were developed using Optiblot ECL kit (Abcam) and visualized via Bio-Rad ChemiDoc MP Imaging system. To restain individual blots for different proteins, blots were stripped with mild stripping buffer (200 mM glycine, 0.1% w/v SDS, 1% Tween 200), 2 x 10 minute washes with shaking, followed by 2 x 10 minute PBS washes with shaking and 2 x 5 minute TBST washes with shaking. After stripping, blots were blocked in 5% milk for 1 hr in preparation for antibody staining as previously described.

### Flow Enabled Tracking of Connexosomes in HEK cells (FETCH)

The FETCH docking assay was completed largely as published^1^. Briefly, replicate multi-well plates were transfected with one connexin construct from each pair being evaluated. Media were changed 16–18 hr post-transfection, and cells from corresponding wells were trypsinized and combined approximately 20 hr post-transfection. The full volume of each combined sample was then replated into fibronectin-coated 96-well plates, resulting in increased cell density and confluency. Following co-plating, samples were incubated for 26 hr, then trypsinized, resuspended in PBS containing 10 U/mL DNase, and fixed with paraformaldehyde. (f/c of 1%). Flow cytometry data was collected on a Cytek Aurora Spectral Cytometer which utilizes the Spectroflo software.

### Fetch Automated Gating Pipeline

FETCH analysis was performed using the automated gating pipeline described previously^23^, with adaptations for spectral flow cytometry data and downstream analysis in R. Each experiment generated FCS files containing fluorescence and scatter measurements for each sample. For this dataset, raw rather than unmixed spectral data were processed, with B1 selected for mEmerald detection and YG3 selected for mCherry detection. The pipeline extracted FSC-A, SSC-A, FSC-H, and the relevant fluorescence channels, then applied sequential gates to identify the analyzed cell population and singlets. Fluorescent events were then displayed on a two-parameter fluorescence plot, with mEmerald and mCherry defining the axes. Quadrant gates were established from the untransfected population, and FETCH scores were calculated as the fraction of fluorescent protein-positive cells that were dual-color positive: FETCH score = Q2/ (Q1 + Q2 + Q3). Gating outputs included scatter plots, density contours, and merged plots with contour overlays for inspection. Flow cytometry-derived FETCH scores were exported to Excel workbooks generated during experimental runs and analyzed in R using the tidyverse framework. Raw data were imported using readxl and standardized using janitor::clean_names() to ensure consistent variable naming. FETCH scores were parsed as numeric values to accommodate mixed-format cells. A quality-control filter flagged runs for manual review when the mEmerald-and mCherry-positive populations differed by more than 2.5-fold or when more than 90% of events were nonfluorescent; low-quality measurements identified during this review were excluded from analysis.

Due to a fluorescence distribution artifact that prevented reliable automated quadrant placement, cytoplasmic FP controls were processed by manual analysis. Single-color controls were used to establish the quadrant gates, with the positivity threshold for each fluorescence axis positioned at the local minimum separating the untransfected and single-color-positive populations. The resulting gates were applied uniformly to all samples within each experiment, and events exceeding both fluorescence thresholds were classified as dual fluorescent.

### Literature Curation of Heterotypic Reference Data

A reference library of 60 papers published studies was compiled to curate reported heterotypic connexin compatibility between specific isoform pairs. Literature assignments were based on evidence from studies assessing gap junction formation and/or functional intercellular coupling, including dye transfer, dual whole-cell electrophysiology, paired-cell recordings, imaging, or biochemical assays. Interactions were classified as positive when published evidence supported docking and formation of conductive heterotypic channels, and negative when published evidence supported failed docking, lack of conductance, or incompatibility. Only interactions with clear consensus were included in diagnostic threshold analysis; interactions with ambiguous or conflicting reports were categorized as conflicted and excluded from threshold derivation.

### Hierarchical clustering and matrix visualization of heterotypic compatibility relationships

Experimentally measured heterotypic FETCH scores for all tested connexin pairs were assembled into a symmetric continuous interaction matrix. To evaluate higher-order organization of heterotypic compatibility behavior, each connexin isoform was represented by its complete vector of heterotypic FETCH interaction scores across all tested partners. Pairwise similarity between connexin compatibility profiles was quantified using correlation distance metrics calculated from continuous FETCH interaction vectors. Hierarchical clustering was performed in SciPy using average linkage, and the resulting dendrogram leaf order was used to arrange both continuous and thresholded heterotypic interaction matrices. Continuous matrices were visualized using matplotlib to examine higher-order compatibility organization. To generate thresholded interaction matrices, experimentally measured heterotypic FETCH scores were benchmarked against previously reported compatible and incompatible connexin interactions curated from the literature. Receiver operating characteristic (ROC) analysis and optimization of Youden’s J statistic were used to identify an empirical FETCH interaction threshold maximizing sensitivity and specificity for interaction classification. FETCH scores above the optimized threshold (0.069) were classified as positive interactions. Positive interactions were additionally annotated by compatibility-class relationship, using previously proposed connexin motif classes to assign each pair as within-group, neighboring-group, or distant/cross-group. Underlying motif-class labels included canonical H-group (Group 2), canonical K-N-group (Group 1), Group 1-other, Group 2-other, and other classifications. Group-level connectivity matrices were then generated by tallying positive interactions between compatibility classes to visualize higher-order organization across the heterotypic interaction landscape. All analyses and visualizations were implemented in Python using pandas, NumPy, SciPy, and matplotlib. Final figure assembly and formatting were performed in Adobe Illustrator.

### EL2 motif sequence analysis

Canonical human connexin protein sequences were obtained from UniProt and aligned relative to previously proposed extracellular loop 2 (EL2) compatibility motif architectures, including the canonical K-N-group (Group 1) motif Φ(K/R)CxxxPCPNxVDCΩΨS and the canonical H-group (Group 2) motif ΦxCxxxPCPHxVDCΩΨS. Reported-length EL2 motif windows were manually extracted relative to the sequence alignment and curated for comparative analysis. Pairwise EL2 motif similarity was quantified using global alignment-based percent identity metrics calculated across the reported-length EL2 motif windows. Pairwise motif similarity values were subsequently compared against experimentally measured heterotypic FETCH scores for all tested connexin pairs. Correlation analyses were performed using Pearson and Spearman correlation coefficients. Representative connexins were selected from the global heterotypic compatibility landscape to illustrate distinct compatibility behaviors identified from clustered continuous and thresholded heterotypic FETCH analyses. All analyses and visualizations were implemented in Python using pandas, NumPy, SciPy, and matplotlib. Final figure assembly and formatting were performed in Adobe Illustrator.

## Statistical information

### FETCH score aggregation and classification

For each connexin pair, well-level FETCH scores were averaged to generate a pair-level mean, and the sample standard deviation (SD) was calculated across wells. For primary interaction classification, pair-level means were rounded to three decimal places. Pairs were classified as FETCH-positive when the rounded mean was greater than or equal to 0.069, the empirically selected threshold determined by ROC analysis using Youden’s J statistic, as described below. The HEK293FT heterotypic dataset included 210 connexin pairs; using this classification rule, 50 pairs were classified as FETCH-positive and 160 as FETCH-negative. All analyses were performed in Python 3.10 using pandas, NumPy, SciPy, Matplotlib, and openpyxl.

### HEK293FT and DKO correlation analyses

For comparisons between HEK293FT and DKO cells, analyses were restricted to connexin pairs with usable pair-level mean FETCH scores in both cell backgrounds. In the homotypic dataset, matched pairs corresponded to the same connexin isoform tested in each background. In the heterotypic datasets, matched pairs corresponded to the same heterotypic connexin combination with valid pair-level mean FETCH scores available in both HEK293FT and DKO cells. Pearson correlation was used to assess linear relationships between FETCH scores, and Spearman correlation was used to assess concordance in score rank order. All correlation p values were two-sided.

For homotypic connexin pairs, 21 matched pairs were analyzed, yielding Pearson’s *r* = 0.85, *p* = 1.29 × 10⁻⁶, and Spearman’s ρ = 0.85, *p* = 1.03 × 10⁻⁶. Across all matched heterotypic connexin pairs, 77 matched pairs were analyzed, yielding Pearson’s *r* = 0.9064, *p* = 8.67 × 10⁻³⁰, and Spearman’s ρ = 0.6910, *p* = 3.51 × 10⁻¹². For Cx46-containing heterotypic pairs, 20 matched pairs were analyzed, yielding Pearson’s *r* = 0.9552, *p* = 5.87 × 10⁻¹¹, and Spearman’s ρ = 0.8541, *p* = 1.65 × 10⁻⁶.

### Literature-based ROC analysis and Youden threshold selection

The threshold analysis was performed using heterotypic connexin pairs with clear literature consensus supporting either positive or negative interaction. Pairs with conflicting reports or no prior literature annotation were excluded. In total, 50 pairs were included in the analysis, comprising 28 literature-positive and 22 literature-negative pairs. The raw, unrounded HEK293FT pair-level mean FETCH score was used as the predictor. Receiver operating characteristic (ROC) analysis yielded an area under the curve (AUC) of 0.8133. The Youden-optimal raw threshold was 0.068931, with Youden’s J = 0.7045, sensitivity = 0.7500, and specificity = 0.9545. This value was rounded to three decimal places to define the final FETCH positivity threshold of 0.069. For comparison, applying a cutoff of 0.069 directly to the unrounded pair-level means yielded a sensitivity of 0.7143 and specificity of 0.9545.

### EL2 sequence-similarity analyses

For each of the 210 heterotypic connexin pairs, pairwise EL2 motif similarity was calculated and compared with the HEK293FT mean FETCH score. Two measures of similarity were evaluated: exact amino acid identity and physicochemical class agreement. For physicochemical class agreement, two amino acids were considered matched if they belonged to the same predefined class: positively charged amino acids (K, R), negatively charged amino acids (D, E), polar amides (N, Q), polar hydroxyl residues (S, T), hydrophobic aliphatic residues (I, L, V, M), aromatic residues (F, Y, W), small nonpolar residues (A), histidine (H), cysteine (C), glycine (G), or proline (P).

Similarity was first calculated across the full 17-amino-acid EL2 motif and then across overlapping 12-amino-acid and 7-amino-acid windows to determine whether smaller EL2 regions showed stronger association with FETCH score. For each window size and similarity metric, the selected window was defined as the window with the largest positive Spearman correlation with the HEK293FT mean FETCH score. If multiple windows produced the same correlation value, the window with the earliest starting position was selected. Raw Pearson and Spearman p values were two-sided.

Because multiple overlapping EL2 windows were tested, we used a permutation approach to assess whether the strongest observed window correlation exceeded what would be expected by chance from the window-selection procedure itself. HEK293FT mean FETCH scores were randomly shuffled among heterotypic pairs to disrupt any true relationship between FETCH score and EL2 similarity. For each shuffled dataset, all candidate windows were rescanned, and the largest positive Spearman correlation was recorded. This procedure was repeated 10,000 times to generate a null distribution of best-window correlations. The selection-adjusted permutation p value was calculated as the fraction of shuffled datasets that produced a best-window correlation greater than or equal to the observed value.

For threshold-based analyses, heterotypic pairs were classified as FETCH-positive or FETCH-negative using the empirically defined FETCH threshold of 0.069, resulting in 50 FETCH-positive and 160 FETCH-negative pairs. For each EL2 similarity measure, similarity distributions between FETCH-positive and FETCH-negative groups were compared using two-sided Mann– Whitney U tests, with rank-biserial effect sizes calculated for group comparisons. ROC analyses were used to evaluate the ability of each EL2 similarity measure to distinguish FETCH-positive from FETCH-negative pairs, and AUC confidence intervals were calculated using stratified nonparametric bootstrap resampling with 10,000 resamples.

Logistic regression was used to test whether full-length 17-amino-acid exact EL2 identity was associated with FETCH-positive classification. The binary outcome was FETCH-positive versus FETCH-negative status, defined using the empirical FETCH threshold of 0.069, and the predictor was pairwise EL2 exact identity scaled from 0 to 1. The model included all 210 heterotypic connexin pairs, comprising 50 FETCH-positive and 160 FETCH-negative pairs. Odds ratios were reported per 10-percentage-point increase in EL2 exact identity, with 95% confidence intervals and two-sided p values.

## Acknowledgements

We thank Dr. Steve Reichow (Oregon Health & Science University) for generously providing the connexin double-knockout cell line; Nicole Renee Brandon (Unified Flow Cytometry Core, University of Pittsburgh) for extensive assistance with flow cytometry analysis and adaptation of data collection and analysis pipelines for spectral flow cytometry; and Dr. Michael Koval (Emory University) for helpful discussions and feedback.

## Author contributions

Conceptualization, E.R.; Methodology, E.R., J.P., and S.Y.; Investigation, S.Y. and A.R.; Formal Analysis, E.R., S.Y., Z.S., and J.P.; Data Curation, Z.S.; Software, J.P. and Z.S.; Visualization, E.R., S.Y., Z.S., and A.R.; Supervision, E.R.; Writing – Original Draft, E.R.; Writing – Review & Editing, all authors.

