## Supplemental Figures 1-9 for "A family-wide atlas of human connexin docking compatibility"

Supplemental Figure 1. Sequential flow cytometry gating strategy for FETCH analysis

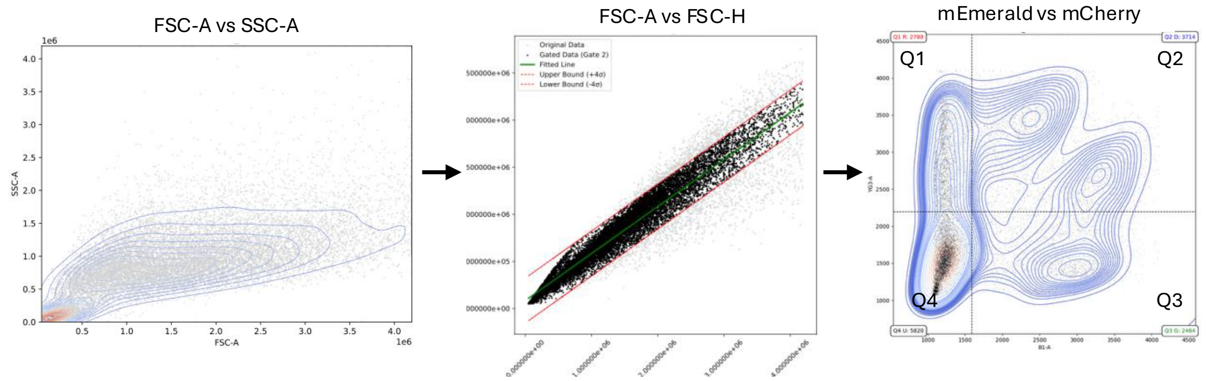


**Supplemental Figure 1. Sequential flow cytometry gating strategy for FETCH analysis.** Representative gating workflow used to quantify FETCH scores from co-cultured connexin-expressing cells. The left panel shows Gate 1, which identifies the presumed HEK293FT cell population and excludes debris based on forward- and side-scatter properties. The middle panel shows Gate 2, which selects singlets from a forward scatter area versus forward scatter height display by retaining events within a defined range of a best-fit line through the singlet population. The right panel shows Gate 3, which evaluates mEmerald and mCherry fluorescence to distinguish single-color and dual-color events. Quadrant gates are set from the untransfected control, with horizontal and vertical thresholds drawn from the contour surrounding the untransfected peak at the 0.6 level. After sequential application of Gates 1-3, FETCH scores were calculated from the fluorescence quadrant gates as Q2/(Q1 + Q2 + Q3), where Q2 represents dual-color events and Q1 and Q3 represent single-color events. This calculation reports dual-color events as a fraction of fluorescent protein-positive events, thereby accounting for variation in the untransfected/background population.

Supplemental Figure 2. Confirmation of connexin expression and relative transfection efficiency across all 21 human isoforms

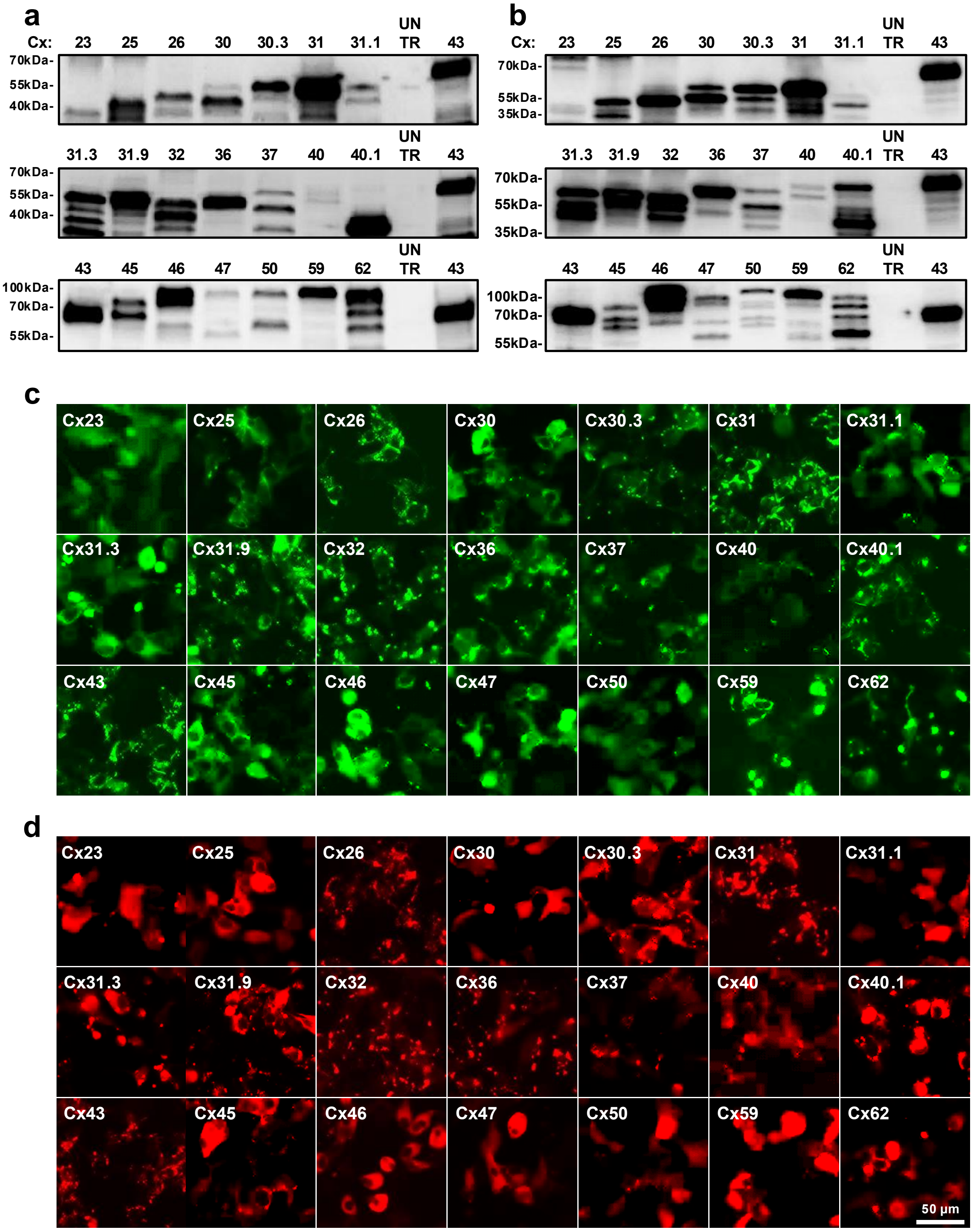


**Supplemental Figure 2. Confirmation of connexin expression and relative transfection efficiency across all 21 human isoforms. (A,B)** Western blot images of HEK293FT cells expressing each of the 21 human connexin isoforms tagged with **(A)** mEmerald or **(B)** mCherry. **(C,D)** Representative widefield fluorescence images of connexin constructs tagged with mEmerald **(C)** or mCherry **(D)**.

Supplemental Figure 3. Alignment and classification of human connexin EL2 motifs

**K-N (Group 1)**

| | |
|---|---|
| Cx25 | KCDLKPCPNTVDCFISK |
| Cx26 | KCNAWPCPNTVDCFVSR |
| Cx30 | KCGIDPCPNLVDCFISR |
| Cx31.1 | KCHADPCPNIVDCFISK |
| Cx32 | KCDVYPCPNTVDCFVSR |
| Cx46 | RCDRWPCPNTVDCFISR |
| Cx50 | RCSRWPCPNVVDCFVSR |
| Cx62 | KCTQPPCPNAVDCFVSR |

**Group 1-Like**

| | |
|---|---|
| Cx31 | CANVAPCPNIVDCYIAR |
| Cx59 | KCHGHPCPNIIDCFVSR |

**H (Group 2)**

| | |
|---|---|
| Cx30.3 | ACSVEPCPHTVDCYISR |
| Cx31.9 | ACAGPPCPHTVDCFVSR |
| Cx40 | VCRRSPCPHPVNCYVSR |
| Cx43 | TCKRDPCPHQVDCFLSR |
| Cx45 | VCSRLPCPHKIDCFISR |
| Cx47 | PCSRQPCPHVVDCFVSR |

**Group 2-Like**

| | |
|---|---|
| Cx31.3 | FACRREPCLGSITCNLS |
| Cx36 | YECNRYPCIKEVECYVS |
| Cx37 | VCQRAPCPYLVDCFVSR |
| Cx40.1 | FPCTRPPCTGVVDCYVS |

**Other**

| | |
|---|---|
| Cx23 | YLCDARSLGENMIIRCM |

**Supplemental Figure 3. Alignment and classification of human connexin EL2 motifs.** Alignment of the reported-length second extracellular loop motif region from all 21 human connexin isoforms, grouped by EL2 motif architecture used for compatibility analyses. Isoforms are organized into K-N, G1-like, H, like, and Other categories. Colored residues highlight conserved and semi-conserved positions across the alignment, including the cysteine/proline-rich EL2 core (orange and green) and positions that vary between the canonical K-N and H motif architectures (pink). This alignment provides the sequence basis for compatibility-group assignments and pairwise EL2 motif similarity calculations across heterotypic connexin pairs.

### Supplemental Figure 4. Representative Homotypic FETCH profiles for all 21 Cx isoforms

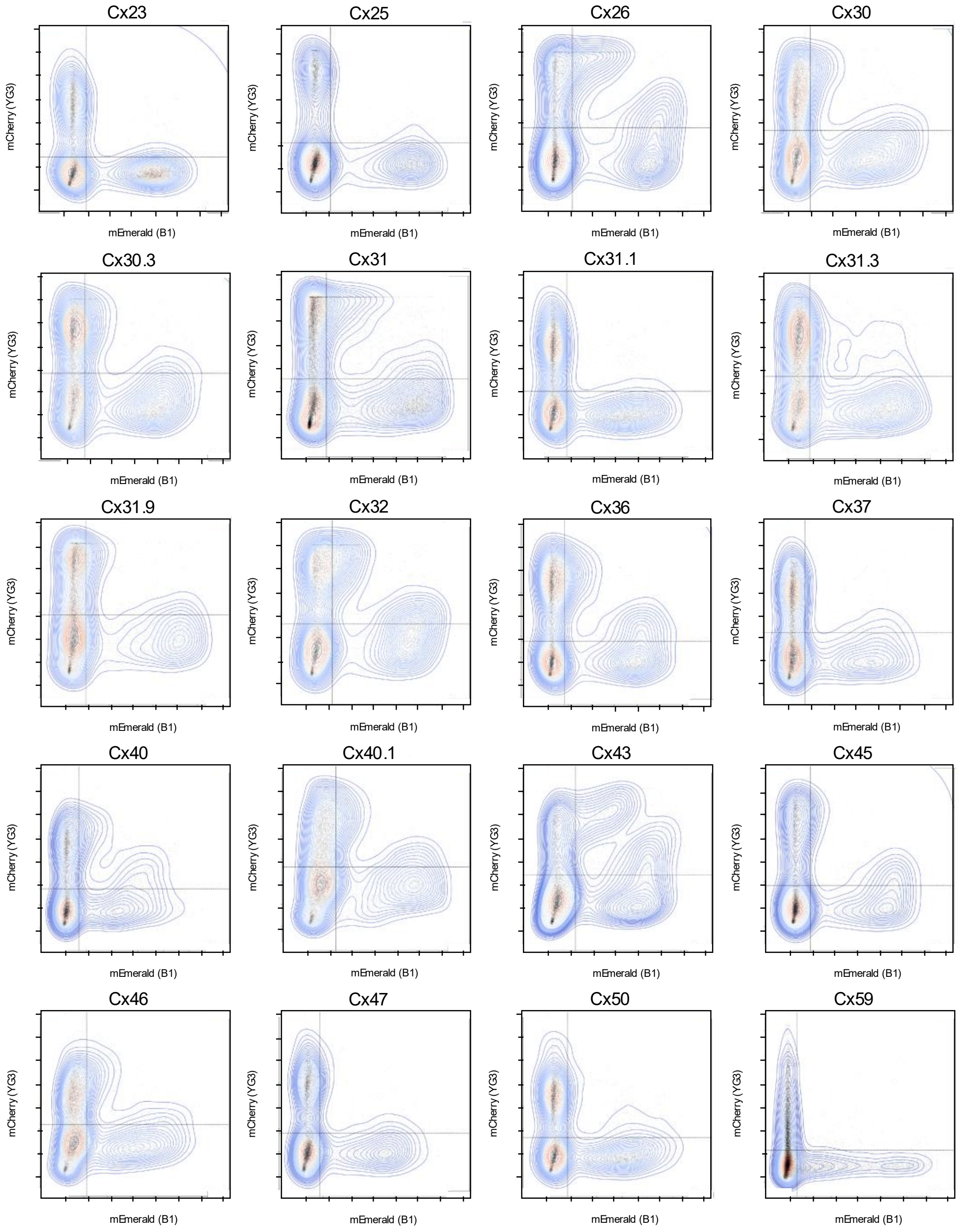


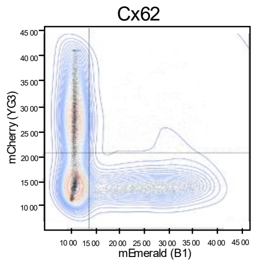


**Supplemental Figure 4. Representative homotypic FETCH profiles for all 21 connexin isoforms.** Representative flow cytometry FETCH profiles obtained from homotypic pairings of each of the 21 human connexin isoforms. Plots illustrate the diversity of dual-color signal distributions observed across the connexin family, reflecting isoform-specific differences in docking-associated transfer and internalization captured by the assay.

Supplemental Figure 5. Homotypic FETCH scores are broadly preserved between HEK 293FT and Cx43/Cx45 double-knockout cells

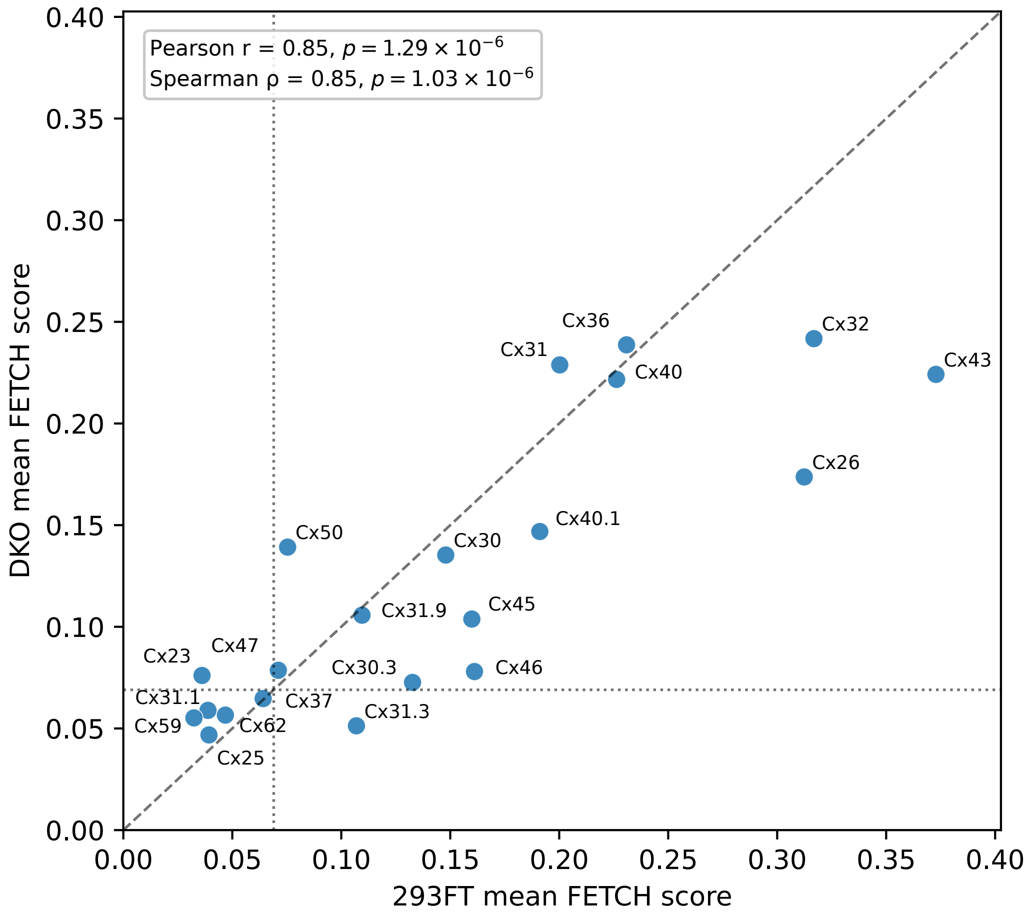


**Supplemental Figure 5. Homotypic FETCH scores are broadly preserved between HEK 293FT and Cx43/Cx45 double-knockout cells.** Comparison of homotypic FETCH scores for human connexin isoforms measured in parental HEK293FT cells and Cx43/Cx45 double-knockout cells. Each isoform was expressed as matched fluorescently tagged constructs and analyzed under the same FETCH workflow used for compatibility measurements.

Supplemental Figure 6. Literature-curated matrix of reported heterotypic connexin compatibility

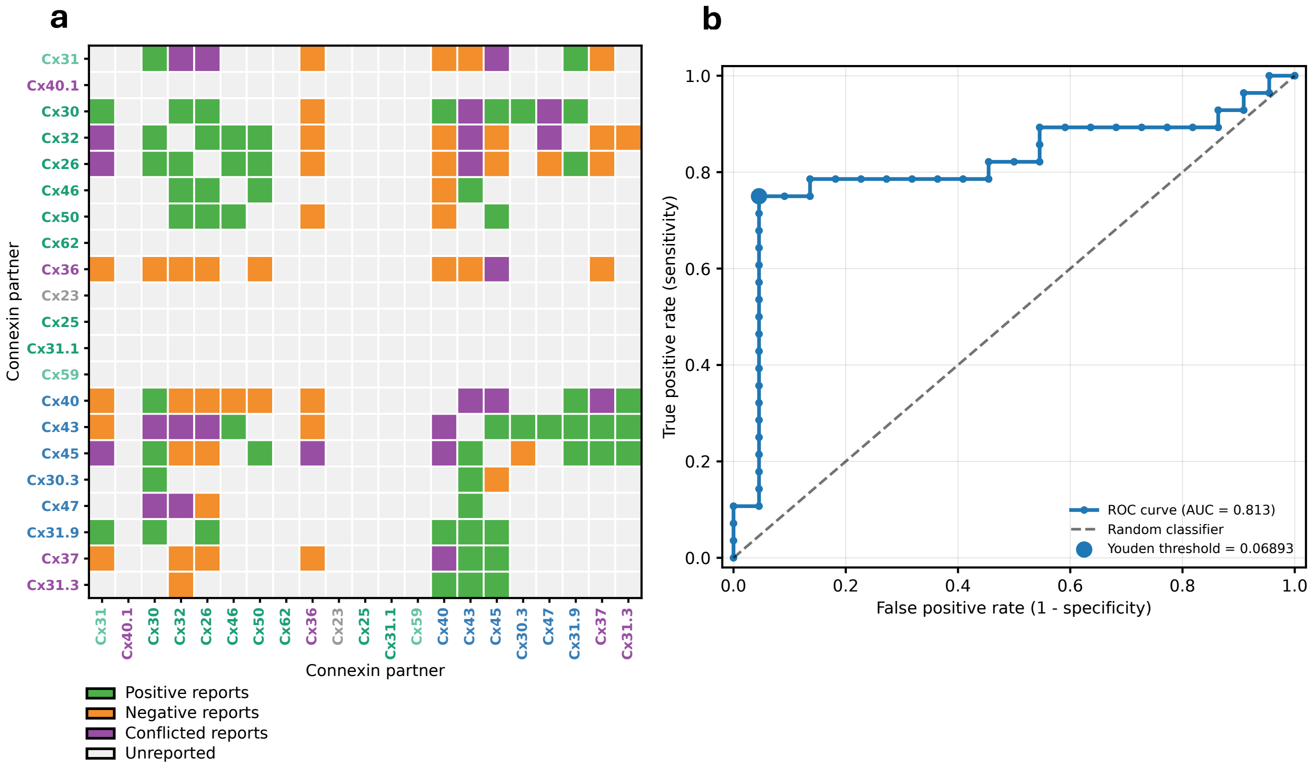


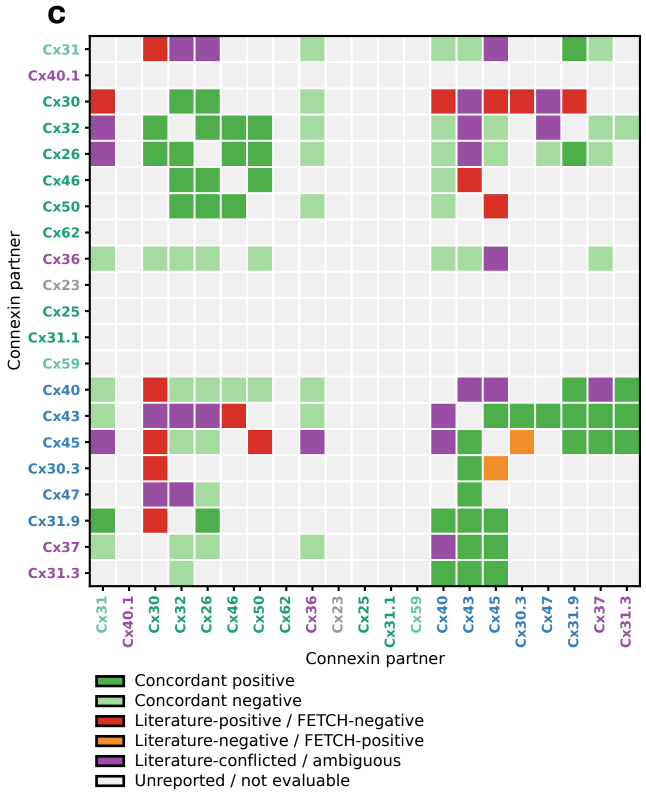


**Supplemental Figure 6. Literature-curated matrix of reported heterotypic connexin compatibility. (A)** Literature-curated matrix of reported heterotypic connexin compatibility across human connexin isoform pairs. Cells indicate whether prior studies reported compatible, incompatible, or conflicting outcomes for each pair. **(B)** ROC analysis used to define the empirical FETCH threshold for high-confidence heterotypic interactions. Continuous FETCH scores were compared against literature-curated compatibility assignments, and Youden's J statistic identified a threshold of 0.069. **(C)** Concordance between FETCH-based classifications and literature-curated compatibility assignments. Pairs are grouped according to whether FETCH and literature classifications were concordant or discordant.

Supplemental Figure 7. Outlier interactions from the global EL2 motif similarity versus FETCH analysis

a

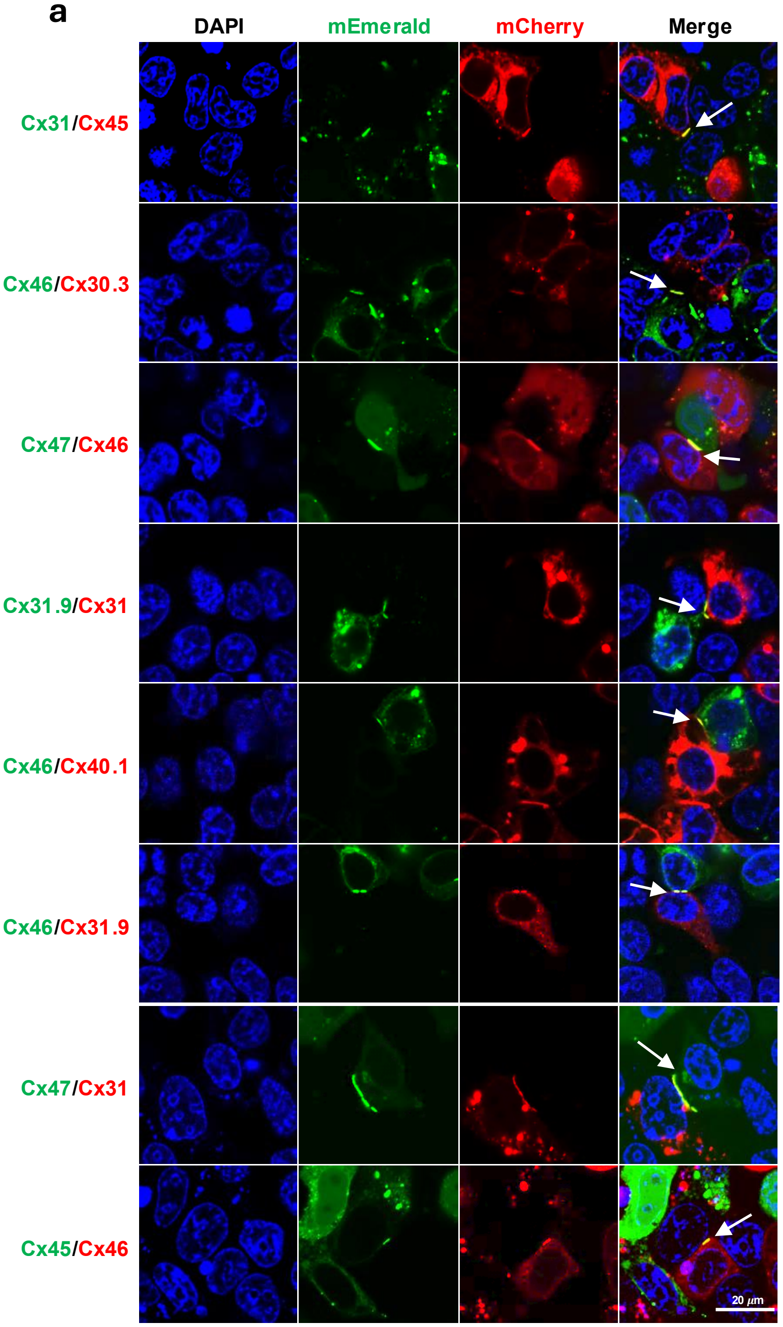


**Supplemental Figure 7. Outlier interactions from the global EL2 motif similarity versus FETCH analysis. (A)** Representative two-color fluorescence images of selected heterotypic connexin pairs identified as outliers in the global EL2 motif similarity versus FETCH analysis. Presumed plaques containing both fluorescently tagged connexin isoforms confirm gap junction formation for the indicated heterotypic pairs. mEmerald- and mCherry-tagged isoforms are indicated by colored labels. **(B)** Summary table of selected outlier pairs, including mean FETCH scores measured in parental HEK293FT cells and Cx43/Cx45 double-knockout cells, as well as percent sequence identity across the reported-length EL2 motif region. Examples are ordered first to show pairs with relatively low EL2 motif similarity but high FETCH scores, followed by pairs with high EL2 motif similarity but low FETCH scores. These examples illustrate that EL2 motif similarity provides useful context for heterotypic compatibility but is not sufficient to fully predict FETCH score across all connexin pairs.

b

| Rank | Pair | 293FTs Ave. | DKOs Ave. | Motif identity |
|---|---|---|---|---|
| 1 | Cx31/Cx45 | 0.2837 | 0.2936 | 41.20% |
| 2 | Cx46/Cx30.3 | 0.2019 | 0.1642 | 64.70% |
| 3 | Cx47/Cx46 | 0.1352 | 0.1327 | 64.70% |
| 4 | Cx31.9/Cx31 | 0.1095 | 0.167 | 41.20% |
| 5 | Cx46/Cx40.1 | 0.1279 | 0.1093 | 11.80% |
| 6 | Cx46/Cx31.9 | 0.116 | 0.1075 | 64.70% |
| 7 | Cx47/Cx31 | 0.1324 | 0.1183 | 41.20% |
| 8 | Cx45/Cx46 | 0.0853 | 0.0489 | 64.70% |

Supplemental Figure 8. Matched comparison of Cx46-containing FETCH scores in HEK 293FT and Cx43/Cx45 double-knockout cells.

a

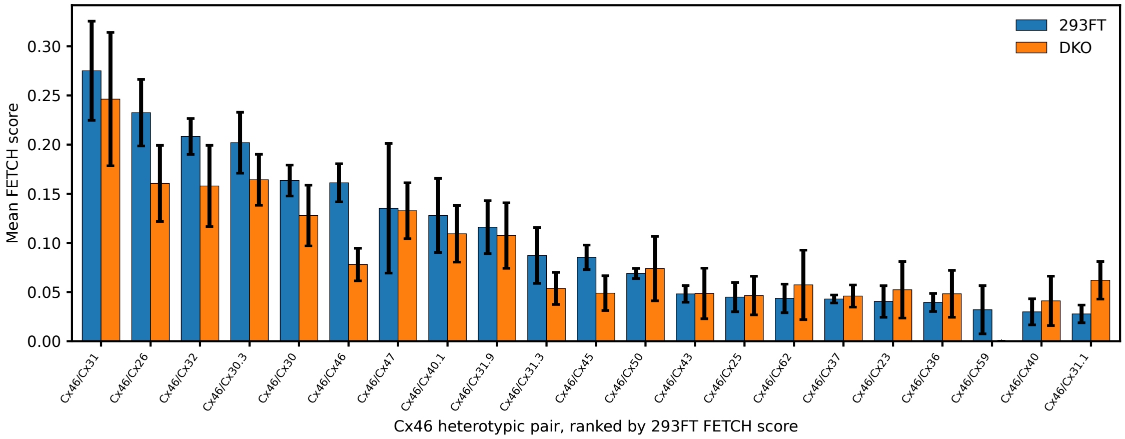


b

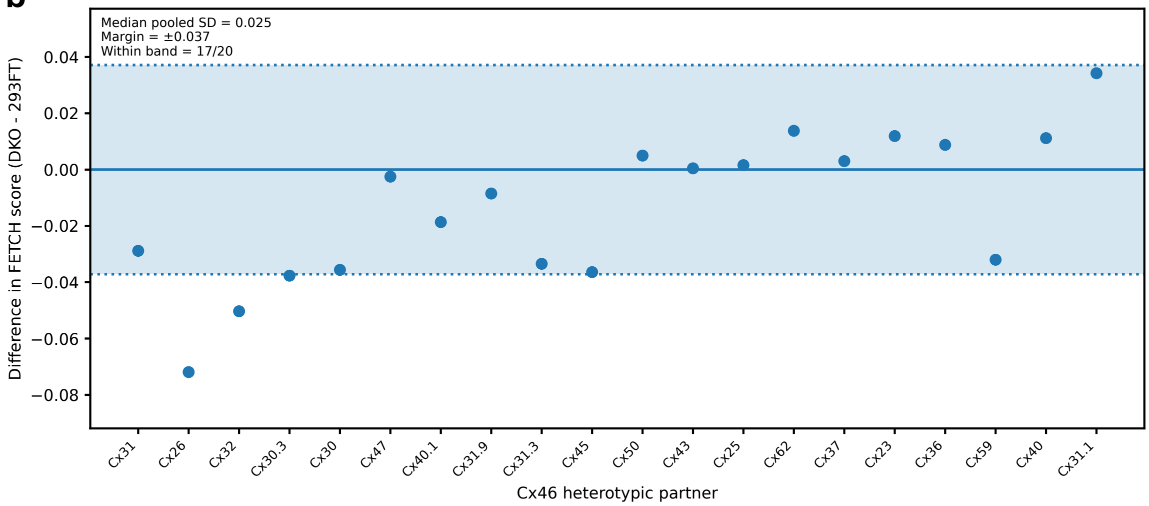


**Supplemental Figure 8. Matched comparison of Cx46-containing FETCH scores in HEK 293FT and Cx43/Cx45 double-knockout cells. (A)** Bar graph comparing Cx46 homotypic and heterotypic FETCH scores measured in parental HEK293FT cells and Cx43/Cx45 double-knockout cells. Cx46-containing pairs are ranked by mean FETCH score in parental HEK293FT cells. Bars show mean FETCH score, with error bars indicating standard deviation across replicates.**(B)** Difference plot showing matched pair-level differences between cell backgrounds for Cx46-containing heterotypic interactions. Each point represents one Cx46 heterotypic pair, plotted as DKO mean FETCH score minus parental HEK293FT mean FETCH score. The horizontal line at zero indicates no difference between cell backgrounds. The shaded equivalence band represents replicate-variability-derived bounds used to define differences considered small relative to assay variation. For each matched Cx46 heterotypic pair, pooled standard deviation was calculated from the parental HEK293FT and Cx43/Cx45 double-knockout replicate standard deviations and replicate numbers. The typical replicate-level variability was then defined as the median pooled standard deviation across Cx46 heterotypic pairs. Equivalence bounds were set as ±1.5 times the median pooled standard deviation, capped at ±0.05 FETCH score units to prevent the tolerance range from exceeding a biologically interpretable difference. In this dataset, the median pooled standard deviation was 0.0279, resulting in an equivalence band of ±0.0418 FETCH score units.

Supplemental Figure 9. Comparison of Cx46 heterotypic and partner homotypic FETCH scores helps distinguish likely positive and negative interactions.

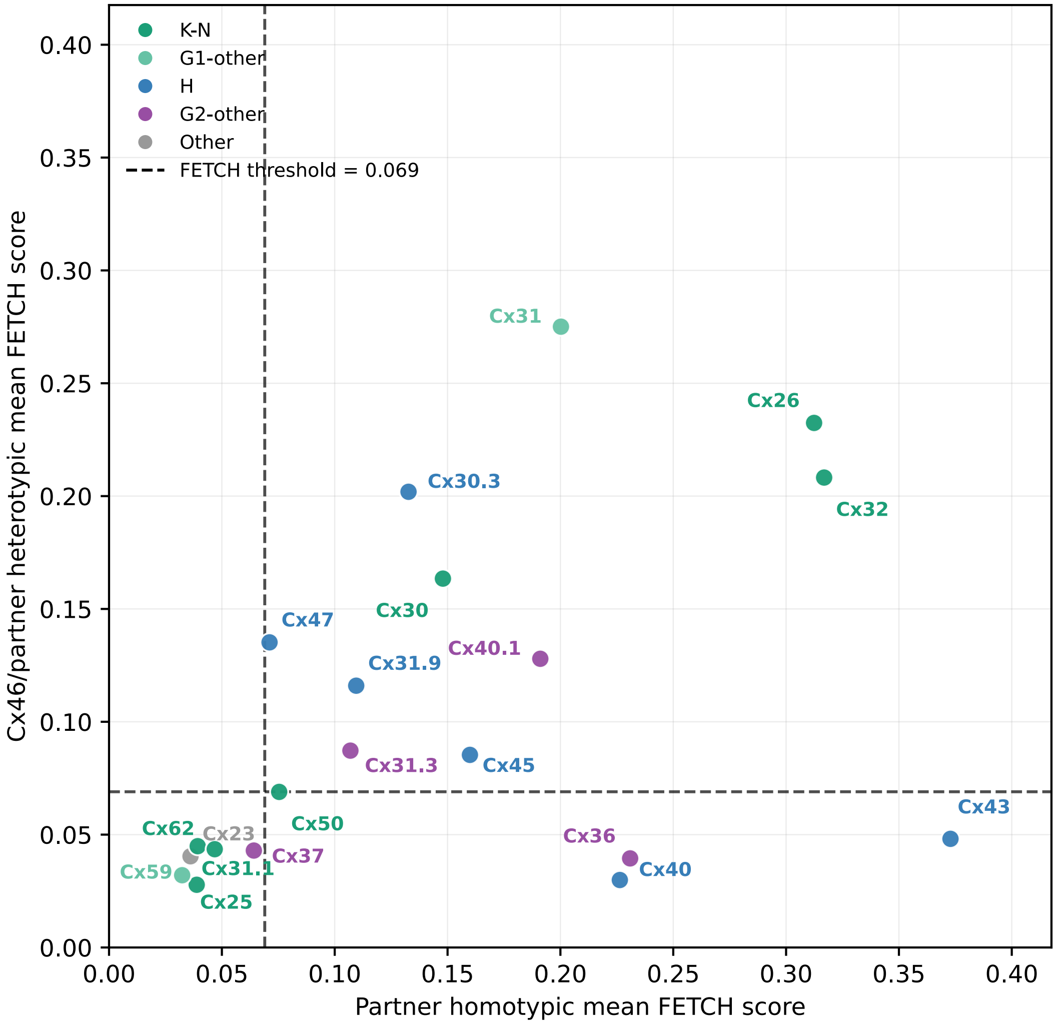


**Supplemental Figure 9. Comparison of Cx46 heterotypic FETCH scores with partner homotypic FETCH activity.** Scatter plot comparing the homotypic FETCH score of each connexin isoform (x-axis) with its heterotypic FETCH score when paired with Cx46 (y-axis). Each point represents one Cx46 heterotypic pairing. Dashed vertical and horizontal lines indicate the empirically defined FETCH interaction threshold of 0.068. Eleven isoforms fell above the threshold in both homotypic and heterotypic measurements. Cx36, Cx40, and Cx43 exhibited above-threshold homotypic scores but below-threshold heterotypic scores with Cx46. Cx23, Cx25, Cx31.1, Cx37, Cx59, and Cx62 fell below the threshold in both measurements. Spearman correlation: ρ = 0.48, *P* = 0.032.
